# Targetable treatment resistance in thyroid cancer with clonal hematopoiesis

**DOI:** 10.1101/2024.10.10.617685

**Authors:** Vera Tiedje, Pablo Sánchez Vela, Julie L. Yang, Brian R. Untch, Laura Boucai, Aaron J. Stonestrom, Alberto Bueno Costa, Sebastià Franch Expósito, Avi Srivastava, Marina Kerpelev, Jillian Greenberg, Mathew Wereski, Amanda Kulick, Kevin Chen, Tianyue Qin, Soo-Yeon Im, Aishwarya Krishnan, Anthony R. Martinez Benitez, Raquel Pluvinet, Merve Sahin, Kamal Menghrajani, Gnana P. Krishnamoorthy, Elisa de Stanchina, Ahmet Zehir, Rahul Satija, Jeffrey Knauf, Robert L Bowman, Manel Esteller, Sean Devlin, Michael F. Berger, Richard P. Koche, James A. Fagin, Ross L Levine

**Author notes:** These authors contributed equally to this work (Vera Tiedje, Pablo Sánchez Vela). These authors jointly supervised this work (James A. Fagin, Ross L Levine).

## Abstract

Anaplastic thyroid cancer (ATC) is a clinically aggressive malignancy with a dismal prognosis. Combined BRAF/MEK inhibition offers significant therapeutic benefit in patients with *BRAF*^V600E^-mutant ATCs. However, relapses are common and overall survival remains poor. Compared with differentiated thyroid cancer, a hallmark of ATCs is significant infiltration with myeloid cells, particularly macrophages. ATCs are most common in the aging population, which also has an increased incidence of *TET2*-mutant clonal hematopoiesis (CH). CH-mutant macrophages have been shown to accelerate CH-associated pathophysiology including atherosclerosis. However, the clinical and mechanistic contribution of CH-mutant clones to solid tumour biology, prognosis and therapeutic response has not been elucidated. Here we show that *TET2*-mutant CH is enriched in the tumour microenvironment of patients with solid tumours and associated with adverse prognosis in ATC patients. We find that *Tet2*-mutant macrophages selectively infiltrate mouse *Braf*^V600E^-mutant ATC and that their overexpression of Tgfβ-family ligands mediates resistance to BRAF/MEK inhibition. Importantly, inhibition of Tgfβ signaling restores sensitivity to MAPK pathway inhibition, opening a path for synergistic strategies to improve outcomes of patients with ATCs and concurrent CH.

## Main

Anaplastic thyroid cancer (ATC) is a relatively uncommon form of thyroid cancer that primarily affects elderly individuals and has a dismal prognosis compared to other thyroid cancer subtypes^1^. The most common truncal oncogenic drivers observed in ATC are activating mutations in the MAPK pathway, primarily *BRAF* and *RAS*^2^. By contrast to differentiated thyroid cancers, ATCs also frequently harbor co-mutations in other genes, most frequently *TP53*^3,4^ and are relatively refractory to standard thyroid cancer therapies including radioactive iodine (RAI), chemotherapy and external-beam radiation. One of the hallmarks of ATCs is their admixture with myeloid cells, primarily macrophages^4–6^. This contrasts with differentiated thyroid cancers (DTC), which are relatively devoid of infiltrating leukocytes^4^. The implications of these changes in the tumour microenvironment (TME) to ATC biology and response to therapy are poorly understood^7^. The recent development of combinatorial treatment with BRAF and MEK inhibitors has led to improved outcomes in class 1 BRAF-mutant ATC patients^8,9^. Although the initial response to MAPK inhibition in ATC is substantive, responses are not durable leading to a median overall survival of around 15 months from diagnosis^10^.

With aging and exposure to genotoxic stressors^11–13^, including RAI^14^, somatic mutations acquired in hematopoietic stem and progenitor cells (HSPCs) display a fitness advantage that leads to their increased representation over non-mutant HSPCs. When this condition is present in the absence of malignant transformation it is referred to as clonal hematopoiesis (CH). CH mutations arise most frequently in epigenetic modifier genes, such as *DNMT3A* and *TET2*^11^. *TET2*-mutant CH is seen most commonly in older patients, and promotes increased HSPC self-renewal, increased myeloid specification^15^ and altered phenotypes in differentiated progeny including mutant macrophages^16^. CH, and especially *TET2*-mutant CH, is associated with an increased risk of atherosclerotic cardiovascular disease and other diseases associated with the aging population^17^.

In a pan-cancer analysis we previously showed that CH, and in particular CH with putative driver mutations (CH-PD), is associated with adverse outcomes in solid tumour patients^12^. However, the mechanisms by which CH can impact solid tumour outcomes, the nature of the specific interactions between CH leukocytes and tumour cells in the TME, and their impact on therapeutic response remain fundamentally uncharted. Recurrent metastatic thyroid cancers, which frequently arise in the aging population, are the solid tumour with the highest CH prevalence^12^. We therefore set out to investigate the impact of CH on clinical outcome and therapeutic response in thyroid cancer, and specifically in ATC.

### Concurrent CH-PD shortens OS in ATC patients

An initial analysis of CH prevalence in 8,810 solid tumour patients in the MSK-IMPACT cohort showed that CH mutations in putative cancer drivers (CH-PD) are associated with decreased overall survival (OS), and the increased risk of mortality was not primarily due to the development of hematological malignancies, but instead to a higher rate of solid tumour progression^12^. In this cohort, thyroid cancer patients showed a particularly high prevalence of CH, most likely due to their advanced age and exposure to iodine radioisotopes^12,14^.

Admixture of CH-mutant hematopoietic cells is frequently observed in next generation sequencing of solid tumour samples, therefore paired blood sequencing is necessary to avoid assigning a CH variant to the tumour cells^18^. Given that CH-mutant leukocytes can potentially interact with epithelial cancer cells within the TME, we leveraged this information to evaluate the tumour infiltrating fraction of CH mutations for thyroid cancer patients with matched genomic analysis of tumor and peripheral blood samples (**Fig 1a**). This analysis revealed that, by contrast to DTC, CH-mutant cells are found at a higher variant allele frequency (VAF) within ATC tumour specimens relative to peripheral blood (**Fig. 1b-c**). The selective clonal enrichment of CH-mutant cells in the TME of ATC relative to blood is also significant when compared to DTC (VAF ratio: 0.14 ATC vs 0.05 DTC, p<0.001) (**Fig. 1c**). These data show that ATCs are characterized by increased clonal representation of CH-mutant cells within their TME.

**Figure 1:**
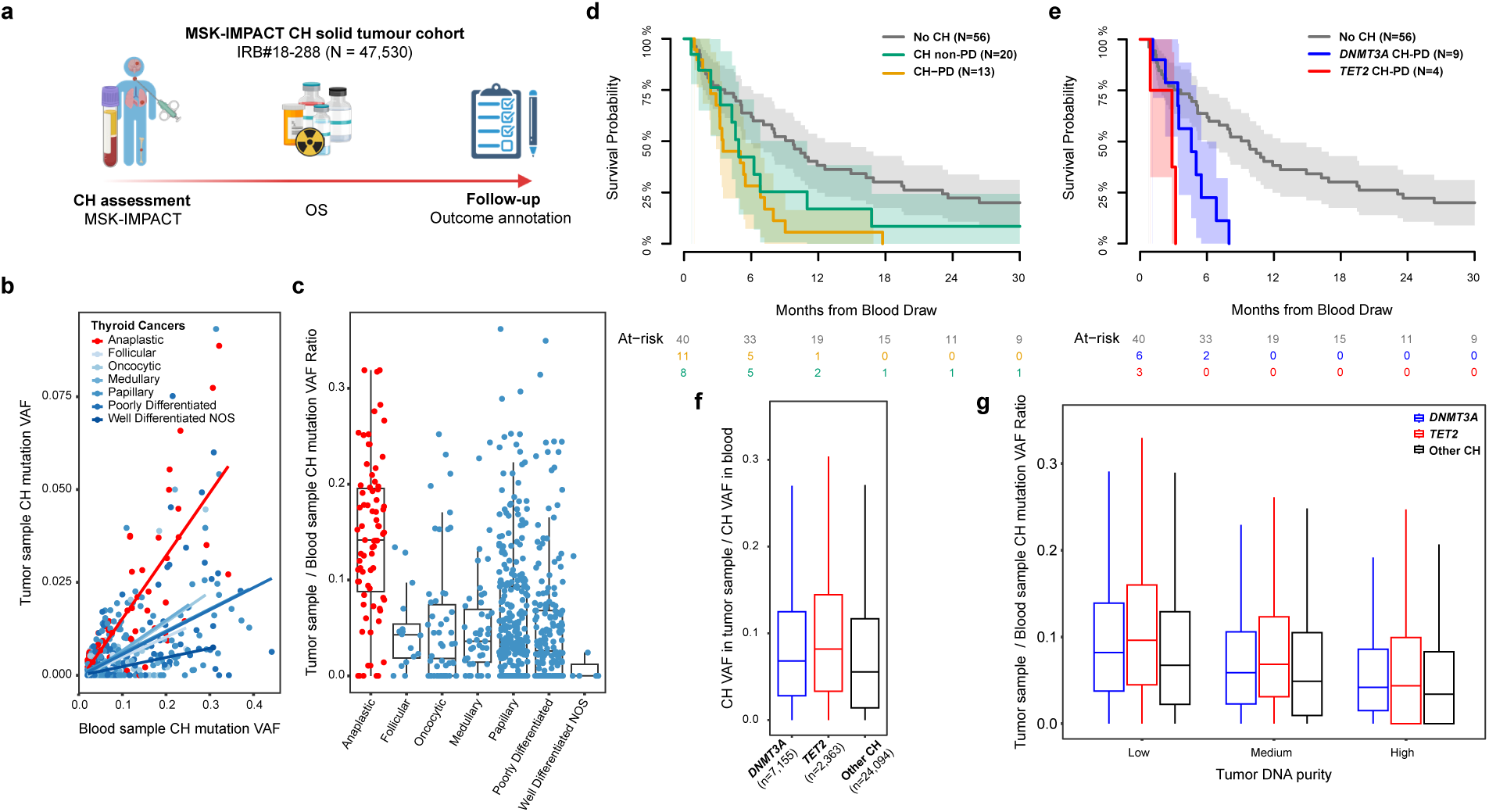
Shorter OS in ATCs enriched with tumor-infiltrating CH-PD mutations. **a**, Patient cohort characteristics; **b**, Relationship between solid-tumour and blood sample CH-VAF among thyroid cancer patients; **c**, Tumour to blood CH VAF ratio for all thyroid cancer subtypes; **d**, Kaplan-Meier curves of ATC patients carrying CH mutations; **e**, Kaplan-Meier curves of ATC patients carrying *DNMT3A* and *TET2* CH-PD mutations; **f**, Assessment of the tumour to blood CH VAF ratio by genotype substratified by tumour purity. ATC: Anaplastic Thyroid Cancer; CH: Clonal Hematopoiesis; CH-PD: Clonal Hematopoiesis in Putative cancer Drivers; HR: Hazard Ratio; OS: Overall Survival; VAF: Variant Allele Frequency; NOS: Not Otherwise Specified.

We next queried a cohort of ATC patients with MSK-IMPACT genomic analysis and curated clinical data **(Extended Table 1)**. In this ATC cohort, age was associated with the presence of CH (p=0.025) (**Extended Table 2**). Having a localized-ATC at the time of MSK-IMPACT testing was associated with improved OS as compared to patients with metastatic disease (HR 0.31, 95%CI 0.18 – 0.52, p<0.001). By contrast, CH-PD detected in the blood of ATC patients was associated with a shorter OS (HR 2.68, 95%CI 1.49 – 4.81, p<0.001). Blood CH variants in non-putative drivers (CH non-PD) had no impact on OS (HR 1.74, 95%CI 0.89 – 3.40, p=0.11) **(Extended Table 3)**. Following correction for age and stage, ATC patients with CH-PD had a significant reduction in OS (HR 2.16, 95%CI 1.12 – 4.14, p=0.021) compared to ATC patients without any CH mutation (**Fig. 1d**) (**Extended Table 4 and 5**). Although the power to detect mutant genotype-specific effects is limited due to the ATC cohort size, the decreased OS signal was also specifically present in *DNMT3A* CH-PD mutation carriers, and even more so in ATC patients with *TET2* CH-PD (**Fig. 1e**).

Given that *TET2* mutations have a greater capacity to promote myeloid lineage bias compared to *DNMT3A* CH mutations^19^, we further evaluated the gene-specific effects of *TET2* mutations on CH clonal tumour infiltration across 47,000 patients in the MSK-IMPACT cohort (**Fig. 1a**). We found increased CH clonal enrichment in solid tumour patients with *TET2*-mutant *CH* as compared to *DNMT3A* and other CH mutations (**Fig. 1d**) among tumour samples with low (mean *TET2* vs *DNMT3A* ratios 0.106 vs 0.093, respectively; p<0.001) and medium tumour purity (mean *TET2* vs *DNMT3A* ratios 0.085 vs 0.072, respectively; p<0.001). This signal was lost in samples with the highest tumour purity (**Fig. 1f**) (mean *TET2* ratio 0.06 vs mean *DNMT3A* ratio 0.058, p=0.1731), likely due to their decreased overall immune cell infiltration. This data shows that CH-PD is common and associated with adverse outcome in ATC. Moreover, CH-clones are selectively enriched within the ATC TME, and *TET2*-mutant clones are most significantly enriched in the TME of solid tumour patients.

### *Tet2*-mutant leukocytes are enriched in ATCs

The observations from the clinical context showing an adverse prognostic impact of CH-PD and *TET2-*mutant CH enrichment in tumours led us to develop a model of concurrent *Tet2*-mutant CH and *Braf^V600E^/p53^−/−^*-mutant ATC (**Fig. 2a**). Our choice of CH-allele was informed by the worse OS conferred by mutations in this gene (**Fig. 1e**), and by the increased tumour/blood ratio of *TET2* CH variants in patients who underwent MSK-IMPACT testing (**Fig. 1b, c**). We transplanted lethally irradiated CD45.1^+^ mice with C57BL/6J CD45.2^+^ bone marrow cells from Wild-type (WT) and mutant cells at a 9:1 ratio (WT:*Tet2*^−/−^) to model the sub-clonal nature of CH. Somatic mutations were generated by tamoxifen-treatment of genetically engineered mouse models (GEMM) harboring a hematopoietic stem and progenitor cell (HSPC)-specific recombinase (*SclCreER^T^*)^20^ and a *TdTomato* reporter^21^ to specifically delete *Tet2*^15^ and activate the TdT reporter in excised/mutant cells (**Fig. 2a**). After engraftment was confirmed, the recipient mice were injected orthotopically with TBP3743 cells, derived from a GEMM of *Braf^V600E^/Tp53^−/−^* ATC^22^ and monitored for signs of tumour progression, including tachypnea, weight loss, and decreased activity. OS following tumour implantation was similar in recipient mice with WT and *Tet2*^−/−^ CH (**Extended Data 2a**), suggesting that tumour growth was not altered by the presence of CH. Multispectral flow cytometric analysis (**Extended Data Fig. 2d**) revealed that although there were no differences in the absolute tumour leukocyte infiltration of the TME (**Extended Data Fig 2b**) or their relative composition (**Extended Data Fig 2c**), *Tet2*^−/−^ CH leukocytes were significantly enriched in the TME (**Fig. 2c**) compared to mutant cell representation in the peripheral blood (**Fig. 2b**). The tumour to blood ratio of mutant cells, which allowed us to correct for variability in bone marrow engraftment for every individual mouse, confirmed 58% higher enrichment of *Tet2^−/−^* cells in the TME compared to the WT counterparts (**Fig. 2d**). Furthermore, evaluation of different organs revealed that the enrichment was primarily present within the thyroid tumours (30% vs 10%) and more limited in non-tumour bearing tissues including lung (4% vs 2%) and liver (4% vs 4%) (**Fig. 2e**).

**Figure 2:**
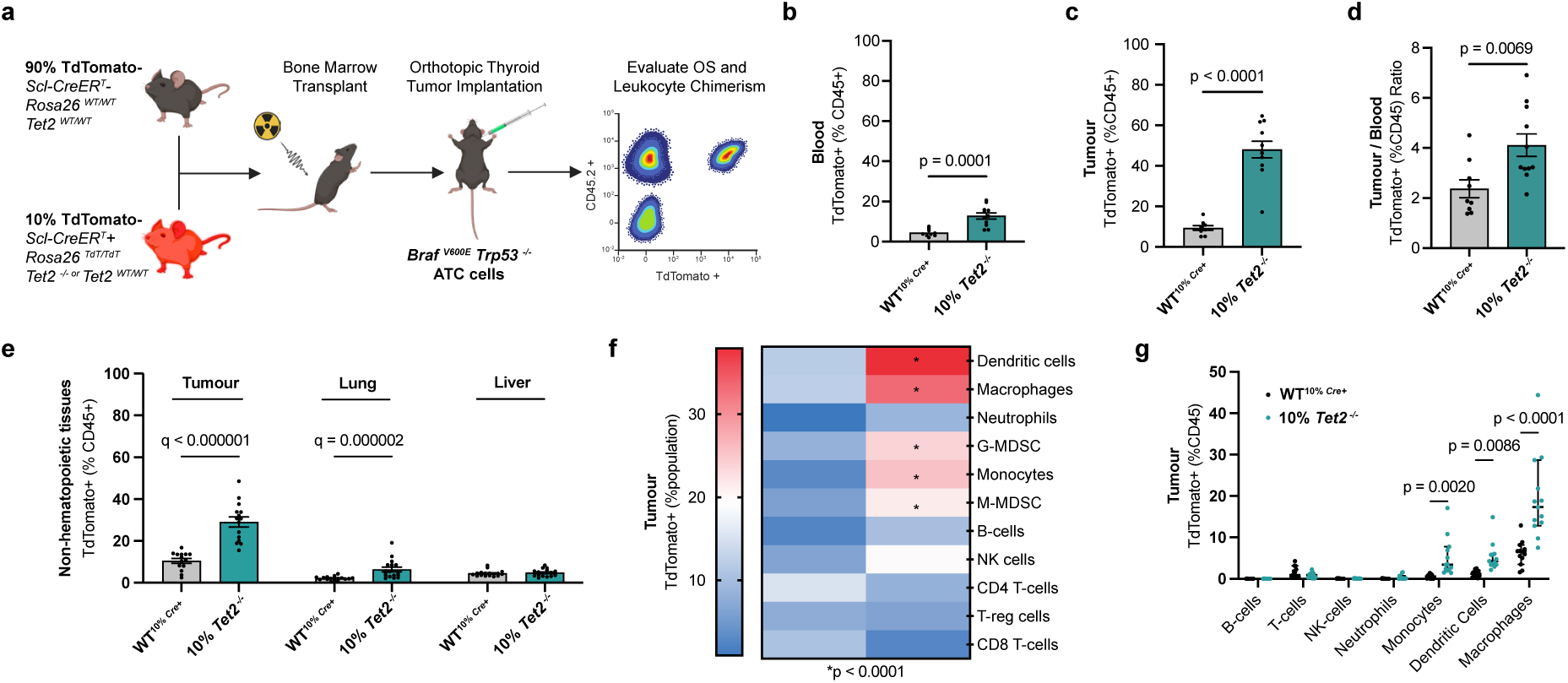
*Tet2*-mutant macrophages accumulate in the tumour microenvironment of ATCs. **a**, Mouse models of Tet2-mutant CH and ATC; **b-c**, Chimerism of total leukocytes (CD45+) in blood (**b**) and ATC tumours (**c**) for WT^10%^ *^Cre+^* (n=9) and 10% *Tet2^−/−^* (n=11) mice, respectively; **d**, tumour/blood ratio for reporter chimerism for CD45+ cells for WT^10%^ *^Cre+^* (n=9) and 10% *Tet2^−/−^* (n=11) mice; **e**, Reporter chimerism of total leukocytes in tumour and non-hematopoietic tissues as indicated from WT^10%^ *^Cre+^*(14 tumours, 17 lung and liver specimens) and 10% *Tet2^−/−^*mice (15 tumours, 17 lung and liver specimens) ; **f**, Chimerism within the respective leukocytic population infiltrating the ATC tumours from 13 WT^10%^ *^Cre+^* and 13 10% *Tet2^−/−^* mice; **g**, Percentage of total leukocytes represented by mutant cells among the leukocytic lineages infiltrating the ATC tumours (n=13 WT^10%^ *^Cre+^* and 10% *Tet2^−/−^* each). Bars show mean with SEM. Two-sided multiple Mann Whitney U tests and Two-stage step-up method of Benjamini, Krieger and Yekutieli for FDR approach (**b-d);** Multiple Mann-Whitney tests (**e**); Two-sided ANOVA with Sidak’s multiple comparisons test, with single pooled variance (**f-g**). CH: Clonal hematopoiesis, ATC: Anaplastic thyroid cancer, WT: wild-type, SEM: standard error of the mean; Dendritic Cells: CD45+ CD11c+ F480-MHCII+; Macrophages: CD45+ CD11b+ Ly6G-Ly6C-F480+; Neutrophils: CD45+ CD11b+ Ly6G+; G-MDSC (Granulocytic-Myeloid Derived Suppressor Cells): CD45+ CD11b+ Ly6G+ Arg1+; Monocytes: CD45+ CD11b+ Ly6C+ Ly6G-; M-MDSC (Monocytic-Myeloid Derived Suppressor Cells): CD45+ CD11b+ Ly6G-Ly6C+ Arg1+; B-cells: CD45+ B220+; NK-cells: CD45+ NK1.1+; CD4 T-cells: CD45+ CD3+ CD4+ FoxP3-; T-reg cells: CD45+ CD4+ FoxP3+; CD8+ T-cells: CD45+ CD3+ CD8+.

To understand the changes in clonal representation, we performed chimerism analysis of each leukocyte population within the tumor to delineate the WT and *Tet2^−/−^* contribution to each hematopoietic subset (**Fig. 2f**). We found that *Tet2*-mutations were enriched in macrophages, dendritic cells, monocytes and both granulocytic and monocytic Arg1+ myeloid-derived suppressor cells (G-and M-MDSCs) infiltrating the tumour (**Fig. 2f**). Furthermore, while there was no overall change in total leukocyte number or individual immune populations infiltrating ATCs (**Extended Data Fig 2 b and c**), lineage subset analysis revealed that *Tet2^−/−^* mutant monocytes (3% vs 0.5%), dendritic cells (4% vs 1%), and especially macrophages (17% vs 6%) (**Fig. 2g**) represented a larger portion of the TME even when their presence in the blood was significantly lower (**Fig 2b**). These results are in line with the relative enrichment of *TET2*-mutant clones infiltrating the TME of solid tumour patients (**Fig. 1b, c and f**) and suggest that blood-based assessments of CH VAF in circulating leukocytes might underestimate the degree of tumour infiltration by CH-mutant leukocytes, and therefore the likely extent of direct CH-epithelial tumour cell interactions.

### CH promotes MAPK inhibitor resistance in ATC

Given that *Tet2^−/−^* CH did not impact tumour growth in the absence of therapeutic perturbation (**Extended Data Fig. 2a**), we hypothesized that the differences in survival observed in patients with concurrent CH (**Fig. 1d and e**) might be mediated by CH-driven resistance to therapy. The current standard of care for *BRAF*-mutant ATCs is treatment with combined BRAF (BRAFi) and MEK inhibitors (MEKi), most commonly dabrafenib and trametinib. We used our mouse model of concurrent *Tet2^−/−^*CH and orthotopically implanted *Braf^V600E^*ATCs to investigate the consequences of mutant myeloid infiltration in the TME on the response to MAPK inhibitor therapy (**Fig. 3a**). Mice with ATC and concurrent CH with either *Tet2* heterozygous (*Tet2^+/−^*) or homozygous (*Tet2^−/−^*) mutations (**Extended Data Fig. 3a**) had significantly worse OS (113 and 68-days median survival, respectively) compared to mice engrafted with only WT hematopoietic cells (147 days follow-up) when treated with BRAFi and MEKi combination therapy (**Fig. 3b**). Whereas BRAF and MEK inhibition induced robust tumour size reduction in mice transplanted with WT bone marrow cells (**Fig. 3c-e**), this treatment was ineffective in mice reconstituted with *Tet2^−/−^* CH (**Fig. 3c-e**). As macrophages were the largest population within the *Tet2-*mutant leukocytes infiltrating the TME (**Fig. 2g**) we hypothesized that they may play a central role in CH-associated ATC resistance to BRAFi/MEKi. Indeed, *in vivo* macrophage depletion via CSF1R signaling blockade using sotuletinib, a CSFR1 inhibitor^23^, in combination with BRAFi and MEKi (**Fig. 3f and Extended Data Fig. 3b**), restored sensitivity to MAPK inhibition in *Tet2*-mutant mice to the levels observed in mice with WT hematopoiesis (**Fig. 3g and Extended Data Fig. 3c**). In summary, this data demonstrated that *Tet2*-mutant CH acts as a driver of primary resistance to MAPK inhibitor treatment in *Braf^V600E^* driven ATCs (**Fig. 3 b-e**), in a macrophage-dependent manner (**Fig. 3g and Extended Data Fig. 3c**).

**Figure 3.**
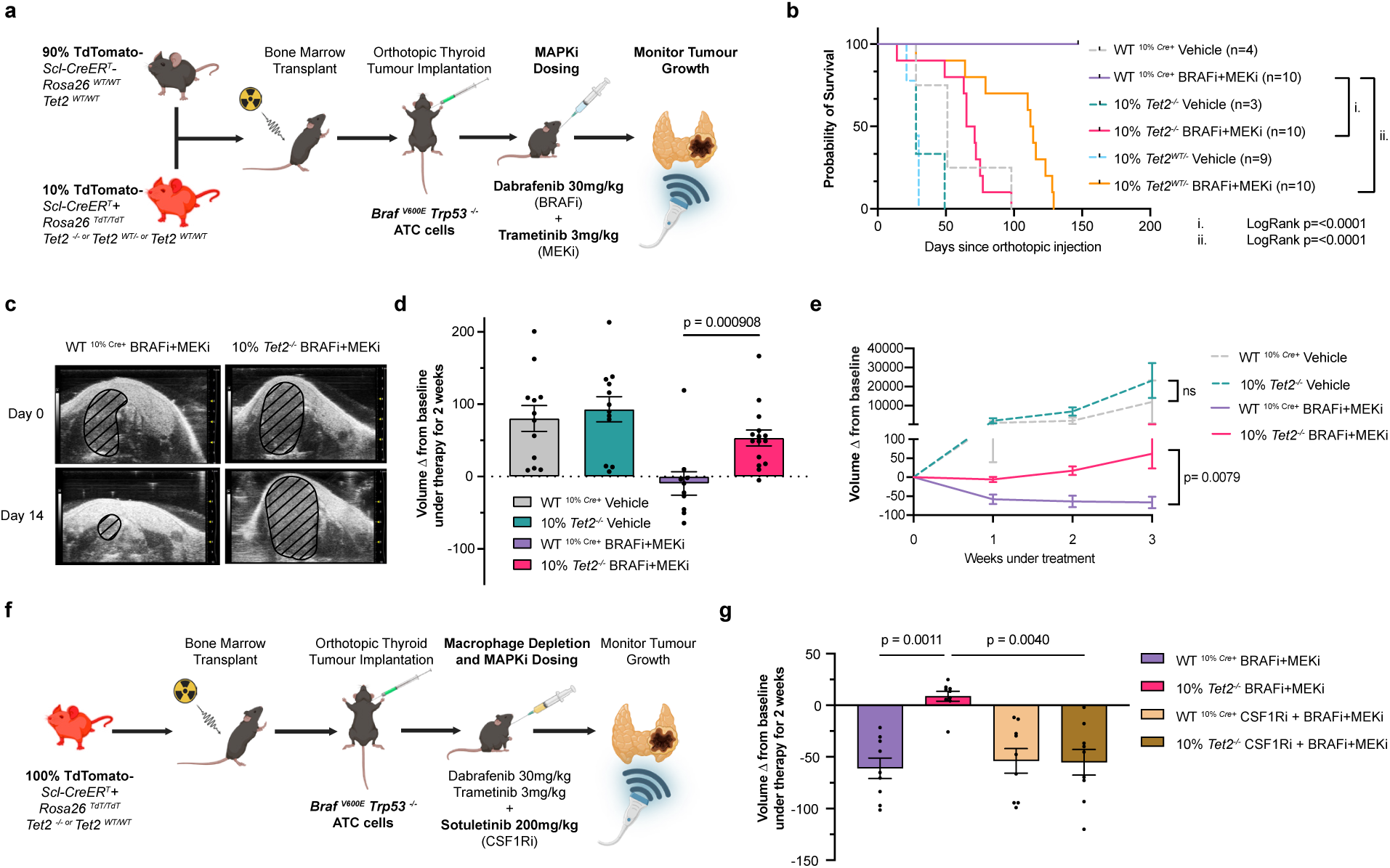
*Braf^V600E^* driven anaplastic thyroid cancers in *Tet2*-mutant CH models are resistant to combined BRAF and MEK inhibition. **a**, Schematic depicting the setup of the pre-clinical trials of combined BRAF and MEK inhibitors; **b**, Kaplan-Meier curves of OS from tumour implantation date until humane-endpoints were met for WT^10%^ *^Cre+^* mice treated with vehicle (n=4) **or** BRAFi+MEKi (n=10), 10% *Tet^+/−^* mice (vehicle (n=9) or BRAFi+MEKi (n=10)) and 10% *Tet2^−/−^ mice* (vehicle (n=3) and BRAFi+MEKi) (n=10) ; **c**, Neck ultrasound images depicting the evolution of ATC tumours under MAPK inhibitor treatment; **d**, Change in tumour volume from baseline for the indicated treatment condition (WT^10%^ *^Cre+^* vehicle n=12, BRAFi+MEKi n=10, 10% *Tet2^−/−^* vehicle n=12 and BRAFi+MEKi n=15) ; **e**, Dynamic changes in tumour volume from the first measure until 3 weeks-post treatment, n=5 mice per indicated genotype and treatment group; **f**, Treatment schema of BRAF+MEK inhibitor and CSF1Ri treatment; **g**, Change in tumour volume from baseline after 2 weeks of treatment (WT BRAFi+MEKi n=9, WT BRAF+MEKi+CSF1Ri n=9, *Tet2^−/−^* BRAFi+MEKi n= 9 and *Tet2^−/−^* BRAF+MEKi+CSF1Ri n=9). Bars show mean with SEM. Log-rank (Mantel-Cox) test (**b**), Two-sided Multiple Mann Whitney U tests and Two-stage step-up method of Benjamini, Krieger and Yekutieli for FDR approach (**d**, **e**, **g**). OS: Overall survival; ATC: Anaplastic thyroid cancer; BRAFi: BRAF inhibitor; MEKi: MEK inhibitor; D0: Day 0 post-treatment; D14: Day 14 post-treatment; CSF1Ri: Colony-stimulating factor receptor 1; SEM: Standard of the mean.

### Alternative TGFβ Signaling Activation in ATC Cells in the context of CH

We next sought to elucidate the mechanisms driving insensitivity of *Braf*-mutant ATCs to BRAF and MEK inhibition in the context of *Tet2^−/−^* CH. We performed CITEseq^24^ of the dissociated tumours 48h post-treatment initiation to capture transcriptional changes in tumour cells and in mutant/WT hematopoietic cells in the setting of the response to BRAFi/MEKi combination therapy (**Extended Data Fig. 4a**). We defined the different compartments in the TME using weighted Nearest Neighbors clustering integrating RNA and ADT-derived data (**Fig. 4a, Extended Data Fig. 4c**), coupled to known lineage-defining transcriptional signatures followed by further manual ADT expression curation **(Extended Data Fig. 4b-e**). To understand the phenotypic changes in tumour cells leading to treatment resistance, we performed pathway score analysis^25,26^ of the ATC cluster (**Fig. 4a**). This revealed that BRAFi and MEKi combination therapy significantly attenuated MAPK transcriptional output in the WT model but did so to a lesser extent in the *Tet2*^−/−^ context (**Fig. 4b-c**)). Further analysis suggested that there was a discordant effect of BRAFi/MEKi therapy on the hypoxia and TGFβ signaling pathways, which were diminished in the WT context but were increased in tumours infiltrated by *Tet2^−/−^*myeloid cells (**Fig. 4c**). Further analysis comparing *Tet2^−/−^* treated vs WT treated ATC tumour cells (**Fig 4c**) showed enrichment for genes involved in the epithelial to mesenchymal transition (EMT) in ATC cells from mice engrafted with *Tet2*-mutant CH, consistent with the previously observed enrichment of the TGFβ pathway, as well as increased expression of mitotic spindle and G2M checkpoint associated genes suggestive of persistent tumour cell proliferation (**Fig. 4d**). We next sought to identify the regulatory networks underlying these transcriptional programs. This led to the identification of the Smad3 transcription factor as the top regulon which was differentially activated in BRAFi/MEKi treated ATC cells in the *Tet2*^−/−^ CH background compared to ATC with WT hematopoiesis (**Fig. 4e**). This finding further underscores the potential importance of the Tgfβ signaling pathway in mediating BRAFi/MEKi resistance *in vivo*. These data, in addition to the cell-cycle state analysis that showed a higher fraction of ATC cells in the G2M-phase in the ATC/*Tet2*^−/−^ CH context compared to ATC cells in the setting of WT hematopoiesis (**Fig. 4f**), support the hypothesis that BRAF/MEK inhibitor resistance may be mediated by activation of alternate pathways driven by *Tet2*-mutant myeloid cells.

**Figure 4.**
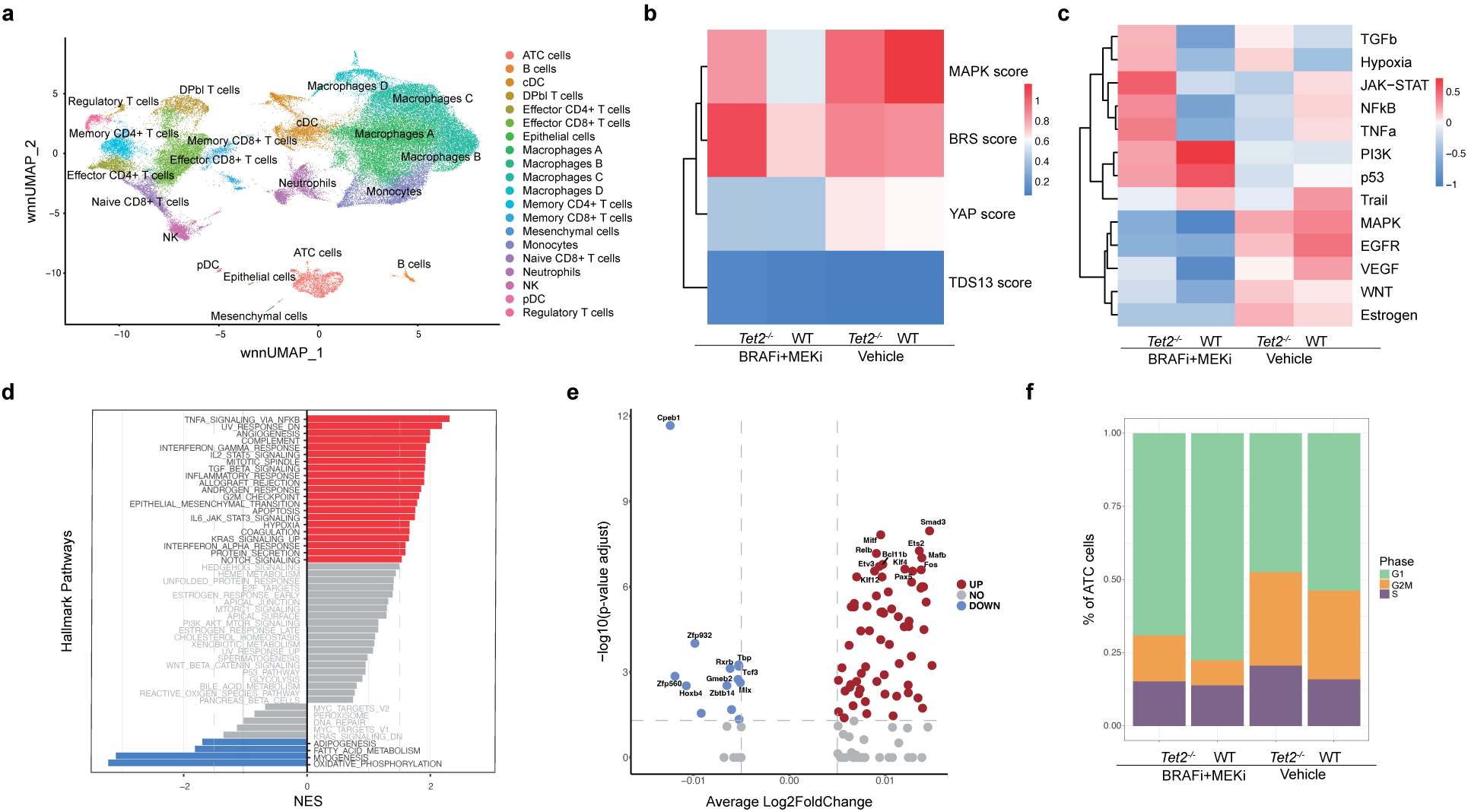
MAPK inhibitor resistant *Braf^V600E^*driven anaplastic thyroid cancers in *Tet2*-mutant CH models show increased proliferation and limited MAPK output. **a,** Weighted nearest neighbors UMAP from the CITEseq analysis of the thyroid tumours 48h post BRAFi+MEKi treatment initiation; **b**, MAPK output, BRS, TDS^25^ and YAP scores^50^ in the ATC cell cluster; **c**, Progeny score analysis of ATC cells; **d**, GSEA analysis of cells in the ATC cluster comparing the *Tet2^−/−^* and WT BRAFi+MEKi treated conditions; **e**, Differential SCENIC analysis of predicted transcription factor-associated gene regulatory network between Tet2^−/−^-and WT BRAFi+MEKi conditions in the ATC cell cluster. The top 10 differential regulons highlighted include: SMAD3, FOS, PAX5, and others significantly upregulated were: STAT3, NFKB1, REL, JUN; **f**, Seurat cell-cycle scoring of the ATC cluster in the CITEseq data. ATC: Anaplastic thyroid cancer; BRS: BRAF-RAS Score; TDS: Thyroid Differentiation Score; GSEA: Gene Set Enrichment Analysis; BRAFi: BRAF-inhibitor; MEKi: MEK-inhibitor; cDC: classic dendritic cells; DPbl T-cells: blasting double-positive T-cells; NK: Natural killer cells; pDC: plasmocytic dendritic cells.

### CH-derived Tgfβ Ligand Secretion Drives Resistance to MAPK inhibitors

We hypothesized that signals emanating from *Tet2*-mutant macrophages promote BRAF/MEK inhibitor resistance (**Fig. 3g**) through secretion of ligands which promote enhanced signaling in the context of ATC-targeted therapy. We therefore performed cell-to-cell communication analysis^27^ with ATC as “receiver” cells and CH mutant vs. WT hematopoietic cells as the “sender” cells. This allowed us to delineate differentially expressed genes (DEGs) into candidate ligand networks that were predicted by the DEG of ATCs in mice reconstituted with *Tet2*^−/−^ versus WT bone marrow cells after BRAF/MEK inhibitor treatment (**Extended Data Fig. 5a**). The highest scoring ligands expressed in CH-mutant hematopoietic cells included Mmp13, Cxcl1, Angpt1, Inhba, Il6, Tgfβ1, Tgfβ3, Il10, Il1b, Ptgs2, Il1a, Ccl5 and Nrg2, among others (**Fig. 5a**). To further prioritize candidate ligands for additional investigation we analyzed their relative expression in the different mutant myeloid cell types admixed within the tumour (**Fig. 5a**). To estimate the relative abundance of a given ligand in the TME, we calculated an effect size score, by multiplying the number of unique UMIs in a given CH-mutant population by the percentage of the population expressing the gene and the fraction of the cell population that composed the TME (**Fig. 5a**). Inhba, Tgfb1, I1b and Ptgs2 were highly expressed by *Tet2*-mutant macrophages. Notably, Mmp13, Inhba, Tgfb1, Tgfb3, and Ptgs2 were differentially overexpressed between the *Tet2*^−/−^ versus WT BRAF and MEK inhibitor treatment conditions in more than one macrophage population (**Fig. 5a**).

**Figure 5.**
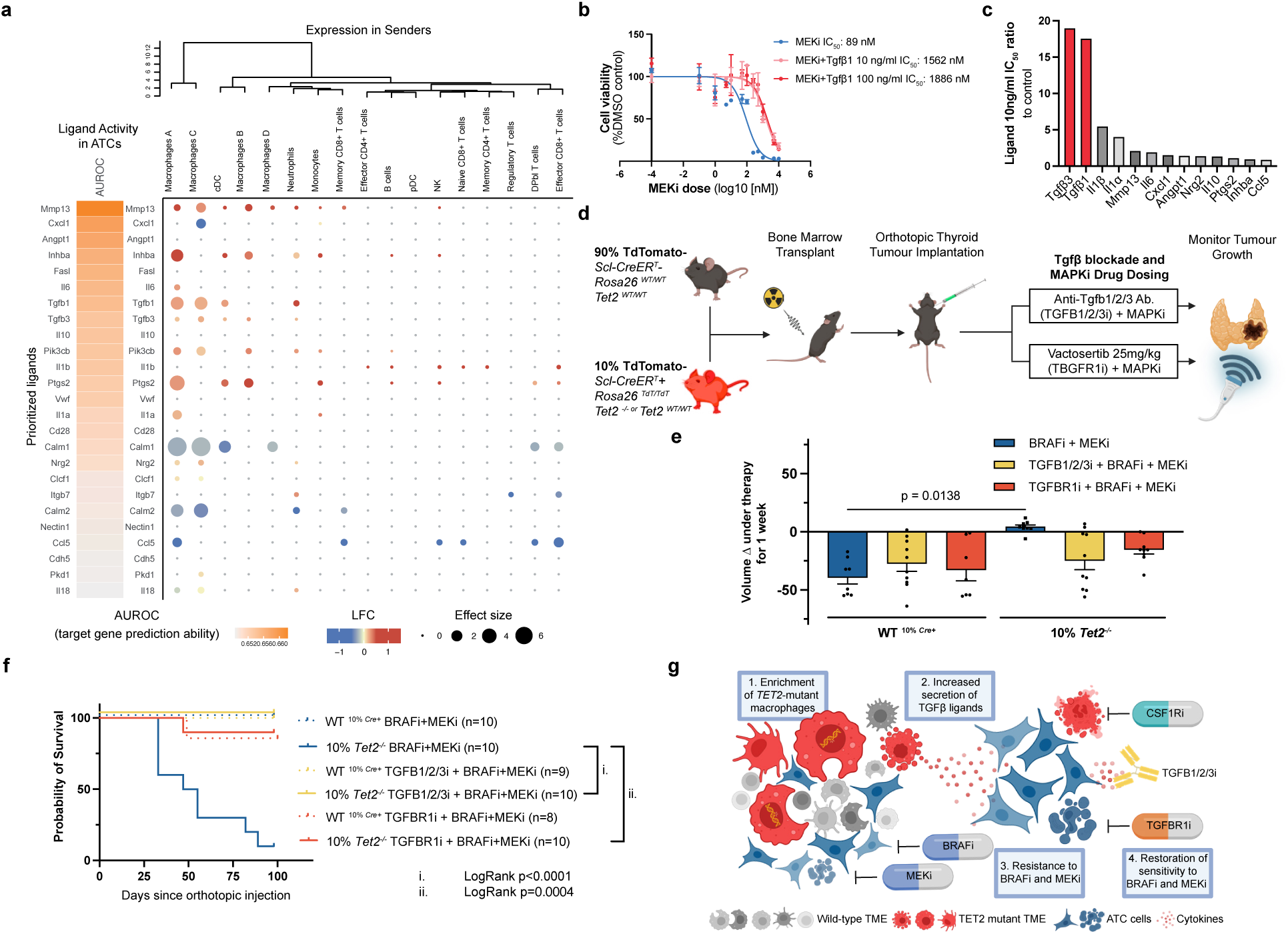
Blockade of *Tet2*-mutant macrophage-derived Tgfβ-signals prevents the development of MAPK inhibitor resistance in the context of *Tet2*-CH. **a,** NicheNet analysis predicting ligands sensed by the ATC cluster based on the DEGs between Tet2 and WT immune infiltrated tumour cells in BRAFi+MEKi treated conditions (yellow left column) and ligand expression by *Tet2*^−/−^ cells (right dot plot); **b**, IC_50_ assessment for trametinib +/− Tgfb1 (10ng/ml or 100ng/ml); **c**, IC_50_ ratio between experimental and control conditions. **d**, Schema of pre-clinical trial with TGFβ pathway inhibitors in combination with BRAFi+MEKi; **e,** Tumour volume after one week of treatment with BRAFi+MEKi (WT ^10%^ *^Cre+^* n=8, 10% *Tet2^−/−^*n=9) +/− TGFBR1i (WT ^10%^ *^Cre+^* n=7 and 10% *Tet2^−/−^* n= 9) or TGFB1/2/3i (WT ^10%^ *^Cre+^* n=10, 10% *Tet2^−/−^* n=10); **f**, OS from tumour implantation date from the BRAFi+MEKi (WT ^10%^ *^Cre+^* n=9, 10% *Tet2^−/−^* n=10) in combination with TGFBR1i (WT ^10%^ *^Cre+^* n=7, 10% *Tet2^−/−^*n=10) or TGFB1/2/3i (WT ^10%^ *^Cre+^* n=10, 10% *Tet2^−/−^* n=10) treated mice; **g**, Schema depicting the enrichment of *Tet2^−/−^* macrophages in ATCs with increased Tgfβ ligand secretion within the TME and the induction of resistance to BRAFi and MEKi that is reversed by blockade of the TGFβ pathway with either the TGFBR1i or TGFB1/2/3i or by depleting macrophages with CSF1Ri. Bars represent mean with SEM, two-sided Multiple Mann Whitney U tests and Two-stage step-up method of Benjamini, Krieger and Yekutieli for FDR approach (**e**); Log-rank (Mantel-Cox) test (**f**). ATC: Anaplastic thyroid cancer; DEG: Differentially expressed genes; BRAFi: BRAF inhibitor; MEKi: MEK inhibitor; TGFBR1i: TGFBR1-inhibitor; TGFB1/2/3i: TGFB1/2/3-inhibitor; WT: Wild-type; TME: tumour microenvironment; CSF1Ri: Colony stimulating factor 1 receptor inhibitor. OS: Overall survival. SEM: standard error of the mean.

Given the selective enrichment of mutant macrophages observed in ATCs (**Fig. 2g**) we hypothesized that ligands secreted by this cell population might drive BRAFi/MEKi resistance (**Fig. 3f, g**). We therefore tested a set of candidate ligands aberrantly expressed in CH-mutant macrophages for their ability to alter the *in vitro* response of TBP3743 ATC spheroids to the MEK inhibitor trametinib (**Fig. 5b-c and Extended Data Fig. 5b-n**). ATC cell viability was decreased by trametinib with an IC_50_ of 89nM. The inhibitory effects of trametinib were markedly attenuated by Tgfβ1(IC_50_ 1562-1886nM) or Tgfβ3 (IC_50_ 533-1688nM) exposure, with other ligands overexpressed by Tet2^−/−^ cells having a more modest effect (**Fig. 5b and Extended Data Fig. 5c**). Tgfβ ligands are known to activate the RAS-RAF-MAPK pathway through recruitment of ShcA to the tyrosine phosphorylated TgfβR1 receptor^28^. Consistent with the impact of pan-Tgfβ activation on MEK inhibitor response, exposure of ATC cells to Tgfβ1 or Tgfβ3 attenuated the inhibition of ERK phosphorylation by the combination of dabrafenib and trametinib (**Extended Data Fig. 5o**).

### Inhibition of Tgfβ Signaling Restores MAPK Inhibitor Sensitivity in ATC with Concurrent *Tet2*-mutant CH

Given that our *in vitro* studies suggested that CH-mutant macrophage Tgfβ secretion could mediate resistance to BRAF/MEK inhibition, we next sought to determine if inhibition of Tgfβ signaling could restore sensitivity of ATC tumours to dabrafenib + trametinib *in vivo* in the context of *Tet2-* mutant CH. We investigated whether treatment with the ALK5 inhibitor (TGFβR1 kinase inhibitor, TGFBR1i) vactosertib^29^ or a pan-Tgfβ ligand-depleting antibody (1D11.16.8, TGFB1/2/3i)^30^ would re-sensitize ATCs to this treatment (**Fig. 5d**). Consistent with our hypothesis, TGFβR1 kinase inhibition and Tgfβ depletion independently restored the ability of the BRAF/MEK inhibitor combination to induce ATC tumour regression in the *Tet2*-mutant CH context (**Fig. 5e**). Of note, treatment with vactosertib or 1D11.16.8 did not impact the efficacy of BRAF/MEK inhibition in ATC cells in the WT hematopoietic cell context, showing that the therapeutic effect was specific to *Tet2*-mutant CH (**Fig. 5e, and Extended Data Fig. 5p and q**). Further supporting the tumour regression data, treatment with either vactosertib or 1D11.16.8 increased survival of mice with concurrent ATC and *Tet2^−/−^*CH in response to BRAF/MEK inhibition, such that their outcome was comparable to treated mice with ATC and WT hematopoiesis (51 days median survival in the dabrafenib-trametinib *Tet2^−/−^* CH group and survival of at least 98 days in both Tgfb-inhibitor-dabrafenib-trametinib *Tet2^−/−^* CH groups) (**Fig. 5e and f**). These data nominate inhibition of Tgfβ signaling as a candidate therapeutic node to restore ATC response to BRAF and MEK inhibition in patients with CH (**Fig. 5g**).

## Discussion

Tumour promoting inflammation has long been recognized as a key pathogenic feature contributing to malignant transformation and adverse prognosis^31^. However, to date this has been challenging to translate into therapeutic benefit in different cancer contexts. In this study we show that *TET2*-mutant CH, which has been shown to increase the risk of hematologic malignancies, cardiovascular disease and other pathogenic states seen with aging can also promote resistance to targeted therapies that inhibit proliferative signaling of *BRAF*-mutant ATCs.

Previous pan-cancer studies have suggested that CH is associated with adverse outcomes in solid tumour patients^12^ and that there could be a causal link between the presence of CH, increased cancer incidence^32–34^, and/or increased tumour agressivity^35,36^. Despite these important insights from molecular epidemiology studies, there is limited insight into the mechanisms by which CH-mutant cells interact with solid tumour cells to mediate adverse outcomes^34,35,37,38^. Most studies of CH in the solid tumour context have focused on patients with advanced and/or high-risk malignancies, given the increased use of genomics as a diagnostic tool in this context. Accordingly, we focused on whether CH-mutant progeny could attenuate the response to tumour-directed therapy and delineated an important link between *Tet2-*mutant CH, Tgfβ signaling and BRAF/MEK inhibitor resistance in ATC. Although ATC is a relatively rare tumour with limited therapeutic options, we hypothesize that additional studies will delineate key mechanistic links between different CH genotypes and reduced response to different anti-cancer therapies in additional cancer subtypes.

Our studies focused exclusively on the contribution of *TET2* mutant CH to adverse prognosis and therapeutic response in ATC. Although *TET2* has been shown to confer a higher risk in the cardiovascular literature than other CH-mutant genotypes^39^, there may be additional ligands/pathways that play a crucial role in how other CH mutant genotypes, such as *DNMT3A*, contribute to adverse prognosis in thyroid cancer and other malignancies. Given the more significant enrichment of *Tet2*-mutant macrophages within the TME in contrast to other CH alleles we hypothesize that the impact of other CH-mutant genotypes on prognosis and therapeutic response may be less substantive and/or require a higher mutant clonal burden to see similar effects. In addition, there may be other myeloid/lymphoid cell types that are the clonal progeny of CH-mutant HSPCs, which contribute to pathogenic heterotypic interactions with tumour cells in ATC and in other cancers. Of note, the pro-interleukin 1β/interleukin 6 secretory phenotype of *Tet2*-mutations in macrophages has been shown to play a key role in mediating heterotypic interactions in non-malignant diseases of aging, including cardiovascular^16,39^, bone^40^, joint^41^, pulmonary^42^, liver^43^, kidney^44^ and infectious disease^45^. In contrast, our data suggests that in ATC, *Tet2*-mutant macrophage secretion of Tgfβ ligands plays a central role in mediating BRAF/MEK inhibitor resistance.

Previous studies in other histologic contexts have suggested that Tgfβ ligands could promote resistance to molecularly targeted therapies^46^; however, this has been limited by the inability to identify a biomarker that enriches for patients most likely to benefit from combination therapies with these drugs. Our work nominates *TET2*-mutant CH as a genetically defined biomarker for adverse prognosis in ATC, which can be specifically targeted through inhibition of aberrant signaling emanating from clonal myeloid progeny derived from mutant stem/progenitor cells. Our work suggests that clonal hematopoiesis, a type of somatic mosaicism, represents a new potential therapeutic node to enhance the efficacy of anti-cancer therapies and may open new opportunities to therapeutically target clonal expansion in a broader spectrum of cancers and age-associated diseases.

## Methods

### Clinical Data

The clinical data from this project has been collected under Memorial Sloan Kettering Cancer Center (MSK) IRB protocol #18-288 (Characterization of Clonal Hematopoiesis in a Cancer Population) using the #12-245 (Genomic Profiling in Cancer Patients) consent form.

Manual ATC clinical data curation was performed retrospectively by annotating this information in a secured institutional database.

### MSK-IMPACT test

The hybridization-capture-based MSK-IMPACT test was performed in solid tumour patients, including those with ATCs as part of their clinical care. As part of this test a tumour and matched peripheral whole blood (PB) genomic DNA (gDNA) samples were sequenced as previously described^47^.

### CH-PD Variant Calling using MSK-IMPACT

As previously described, CH variants were ascertained from the peripheral blood sample first and were further annotated as contaminating the tumour sample if the same variant detected in the blood was found infiltrating the tumour^18^. CH-PD variant annotation was performed as previously described^13^. Tumor content in tissues on MSK-IMPACT were estimated using FACETS as previously described^48^.

### Mouse Models of CH

To generate the genetically engineered mouse models (GEMM) of CH, the previously generated C57BL6/J GEMM of *Tet2^fl/fl^* ^15^ were crossed with inducible *HSC-SCL-CreER^T^* ^20^ and a *TdTomato* reporter^21^. To induce the expression of the *CreER^T^*4–16-week female mice received three doses of Tamoxifen (**Supplementary Table 1**).

To model CH, which frequently occurs as a somatic subclonal event, CD45.2 or CD45.1^49^ recipient female mice between 6 and 16 weeks of age were myeloablated upon a single 978 cGy exposure to ^137^Cs inside a Gammacell irradiator. Eighteen to 24 hours after irradiation, the mice were injected via the tail vein (TVI) with 0.2ml of sterile PBS containing 2×10^6^ red blood cell (RBC) lysed whole bone marrow cells from GEMM donors at a 1:10 Mutant:WT ratio^16^.

### Mouse Models of ATC

To model ATC we used the TBP3743 cell line derived from ATCs generated from a TPO-CreER^T^ *Braf^V600E/WT^*and *Trp53^−/−^* GEMM^22^. At least one-month post-BMT, the C57BL6/J mice were injected with 5ml of sterile PBS containing 5 × 10^3^ TBP3743 cells into the right thyroid lobe under ultrasound-guidance (F2 micro-Ultrasound, Visualsonics; **Supplementary Table 2**) using a 30Gx1/2 needle (**Supplementary Table 3**) under isoflurane anesthesia.

To monitor tumour growth, mice were ultra-sounded on a weekly basis under isoflurane anesthesia using a Vevo F2 micro-Ultrasound. To calculate the tumour volume, a series of 45 images were taken, and the tumour area was manually drawn. An elliptical figure was reconstructed to estimate the volume. The delta was calculated as follows: New volume measurement/Baseline measurement volume. To monitor the overall health condition and evaluate when a humane endpoint had been reached, mice were monitored for early signs of tachypnea, weight loss and overall decrease in activity.

### Drugs

To prepare Tamoxifen and Sotuletinib (BLZ945, CSF1R inhibitor), the respective powder was diluted in corn oil and mixed overnight at 37C (**Supplementary Table 1**).

To prepare dabrafenib and trametinib, the powder was resuspended in 2% DMSO added to 0.5 % HPMC, 0.2% Tween80 in H2O pH8.0. Vactosertib (TGFBR1 inhibitor, EW7197) was prepared in DMSO and artificial gastric fluid (95 mM HCl, 38 mM NaCl and 3.6 mg/mL pepsin). The respective solutions were mixed overnight and kept at 4C for up to a week (**Supplementary Table 1**). To prepare the antibodies, the solutions were diluted in their specific solvents (**Supplementary Table 1**).

### Cell line generation

The *Braf^V600E^ Trp53^−/−^* ATC cell line TBP3743 was a kind gift from Dr. Sareh Parangi^22^ The cell line was grown in DME-HG medium supplemented with 5% of fetal bovine serum (FBS) and 1% of penicillin/streptomycin/L-glutamine (PSG) (**Supplementary Table 3**) and maintained at 37°C and 5% CO2 in a humidified atmosphere.

### Biospecimen processing

Around 30ul of blood was collected from the submandibular vein into EDTA coated tubes. Before downstream processing, red blood cells (RBC) were lysed. To isolate tumour and other solid organs, mice were euthanized in a CO_2_ chamber followed by cervical dislocation before the organs were dissected using clean surgical tools. To generate single-cell suspensions from the solid organs and thyroid tumours the freshly isolated tissue was incubated at 37C with collagenase and DNAse for 45 minutes. The single cell suspension was later quenched with PBS + 2% FBS and filtered through a 70mm mesh (Materials are listed in **Supplementary Table 3**).

### Flow Cytometry

Single cell suspensions were incubated with a viability dye in PBS, followed by Fc block and the extracellular antibody cocktail in brilliant stain buffer for at least 15 minutes each. After quenching the staining reaction with PBS + 2% FBS, cells were fixed and permeabilized by incubation with the Foxp3 / Transcription Factor Staining Buffer Kit (Tonbo Biosciences, **Supplementary Table 3**) for 1h and subsequently stained with intracellular antibodies. The antibodies and reagents are listed in **Supplementary Table 3 and 4**. The analysis was done with Cytek Aurora (**Supplementary Table 2**) and FloJo Version 10.7.2. The full gating strategy is provided in **Extended Data Fig. 2d**.

### CITEseq sample preparation

To generate the CITEseq data (**Extended Data Fig. 4a**) and obtain enough cells for sequencing, non-competitive bone marrow transplants with WT^Cre+^ or *Tet2^−/−^* bone marrow were performed. After engraftment and tumour injection, tumours were collected 48h post-treatment initiation with BRAF/MEK inhibitors, dissociated, and stained with TotalSeq-B (**Supplementary Table 5**), and DAPI. TdTomato positive (TDT+) and TdTomato negative (TDT-) DAPI negative cells were sorted using a Sony SH800. Finally, every population was stained (hashed) using TotalSeq-B hashing antibodies (**Supplementary Table 5**).

### CITEseq (transcriptome sequencing)

Dissociated tumour cells were stained with Trypan blue and Countess II Automated Cell Counter (ThermoFisher) was used to assess both cell number and viability. Following QC, the single cell suspension was loaded onto Chromium Next GEM Chip G (10X Genomics PN 1000120) and GEM generation, cDNA synthesis, cDNA amplification, and library preparation of 15,000 cells proceeded using the Chromium Next GEM Single Cell 3’ Kit v3.1 (10X Genomics PN 1000268) according to the manufacturer’s protocol. cDNA amplification included 11 cycles, and 64-136 ng of the material was used to prepare sequencing libraries with 12 cycles of PCR. Indexed libraries were pooled equimolar and sequenced on a NovaSeq 6000 in a PE28/88 run using the NovaSeq 6000 S4 Reagent Kit (200 cycles) (Illumina). An average of 308 million paired reads was generated per sample.

### CITEseq (cell surface protein feature barcode sequencing)

Amplification products generated using the methods described above included both cDNA and feature barcodes tagged with cell barcodes and unique molecular identifiers. Smaller feature barcode fragments were separated from longer amplified cDNA using a 0.6X cleanup with aMPure XP beads (Beckman Coulter catalog # A63882). Libraries were constructed using the 3’ Feature Barcode Kit (10X Genomics PN 1000276) according to the manufacturer’s protocol with 10 cycles of PCR. Indexed libraries were pooled equimolar and sequenced on a NovaSeq 6000 in a PE28/88 run using the NovaSeq 6000 S4 Reagent Kit (200 cycles) (Illumina). An average of 88 million paired reads was generated per sample.

### CITEseq analysis

Sequencing reads from the mRNA library were aligned to the mm10 reference genome using Cell Ranger software (version 7.0.0). Sequencing reads from the antibody derived tags (ADTs) and hashtag oligonucleotides (HTOs) libraries were processed using Alevin-Salmon (version 1.6.0) and ADTs and HTOs were collectively tabulated into a count matrix. Count matrices from Cell Ranger and Alevin-Salmon were then used as input into Seurat package (version 4.3.0) in R (version 4.2.0) for all downstream analyses (**Extended Data Fig. 4b**).

Taking in the raw and filtered matrices from Cell Ranger as input, the SoupX algorithm (version 1.6.2) using default parameters was used to remove the ambient mRNA count which the algorithm down-weights using regression. The HTO Demux algorithm from the Seurat package was utilized to demultiplex the single cells expressing the HTOs into *Tet2^−/−^* CH TDT+ and *Tet2^+/+^*CH TDT-cell populations within the TME with a negative binomial distribution background count model. The cells were filtered for quality control. Genes with three or more cells supporting their expression were kept. Cells with more than 200 mRNA features and more than 500 UMI counts were kept. The cells with <= 10% mitochondrial genes and >= 5% percent ribosomal genes were kept for downstream analysis. Following the filtering, a total of 22,990 genes along with 16 cell surface proteins in 60,875 single cells were included in the final analysis. The full single cell data set consists of four pooled samples of {*Tet2^−/−^*, WT} and {DT, VEH} conditions.

To visualize cells based on an unsupervised transcriptomic analysis, unsupervised clustering in principal component analysis (PCA) was conducted by using the top 2,000 highly variable genes from all the samples. Cell cycle scores were calculated using the CellCycleScoring function in Seurat version 4.3.0 with default parameters. For the RNA assay log normalization and KNN clustering was performed with number of pcs = 30 and res = 0.8. For the ADT assay centered log ratio (CLR) transformation was performed and the data was KNN clustered with number of pcs=15 and res=0.5. For the dimensionality reduction and RNA and ADT data integration, the weighted nearest neighbor algorithm was used to produce an integrated WNN UMAP of RNA and ADT data (**Extended Data Fig. 4c**).

The scType algorithm (version 1.0.1) was used in combination with Haemopedia Mouse RNAseq, Immgen, Cell marker, and Panglou marker databases to annotate cell types of 60,875 single cells (**Supplementary Table 11**). ADTs for each cell population were then manually curated and used to make confident decision calls for the cell typing of single cell populations. Heatmaps were generated for each subpopulation (**Extended Data Fig. 4d-e**). The FindMarkers function in Seurat was used to perform differential gene expression analysis on the pairwise contrasts between the four conditions using a non-parametric Wilcoxon rank sum statistic and significant genes were called at an FDR and average log fold change thresholds of (FDR ≤ 0.05 and avglog2FC >= 0.25). A comprehensive long list of differentially expressed genes were obtained by setting the log2FC in the FindMarkers function to minimal threshold (log2FC < 0.00001) and Genes Set Enrichment Analysis (GSEA) of the Hallmark pathways was performed on these differentially expressed genes that were ranked their by avglog2FC. The top ranked Hallmark pathways were then visualized with a NES enrichment waterfall plot with thresholds (FDR ≤ 0.05 and NES >= 1.5). Progeny analysis was performed to visualize the pathways activated in the cell populations and progeny scores were plotted in heatmaps with hierarchical clustering using the complete linkage and Pearson correlation metric in the pheatmap package (v1.0.12).

ATC scores (MAPK, BRS, YAP, TDS13 scores) were calculated for ATC cells for the four samples {*Tet2^−/−^*, WT} and {DT, VEH} conditions. A list of mouse marker genes for each score was provided with a directionality (−1, 1) (**Supplementary Table 7-10**). ATC scores were calculated by taking the average expression of each marker gene in the ATC cell population and multiplying the genes’ average expression by their directionality (−1,1), taking the sum of the result, and normalizing by the total number of gene members in the whole markers set. Fagin scores were visualized with hierarchical clustering with the standard complete linkage and Pearson correlation using pheatmap package (v1.0.12).

Regulon analysis was performed with pySCENIC (version 0.12.1) and R SCENIC package (version 1.3.1). The Tet2 DT condition was contrasted against WT DT and the AUC scores were differentially assessed using a non-parametric Wilcoxon rank sum statistic. Cell-to-cell communication analysis was conducted using the NicheNet algorithm (version 2.0.0). The ATC cells were set as the ‘receivers’ and the TDT+ infiltrating immune cell populations were set as the ‘senders’. The ligands were ranked by the ligand activity of the cells as measured by AUROC predicting the gene expression in the receivers. The thresholds for the differential expression for the NicheNet algorithm were set to nominal p-value <0.05 and logFC >0.25 with min.pct >0.10.

### 3D Spheroids and IC_50_ Assays

For the 3D spheroid assays, 500 TBP3743 cells were plated in Ultra-Low Adherence 96 well plates and cultured with increasing concentration of trametinib plus/minus cytokines for 5 days in 5% FBS, 1% Penicillin/Streptomycin/Glutamine and 0.5% methylcellulose. After this time, the spheroids were incubated with CellTiter-Glo® 3D Cell Viability Assay reagents and quantified in a Promega GloMax® 96 Microplate Luminometer (**Supplementary Table 2**). Absolute viability values were converted to percentage viability compared to DMSO control values of the respective cytokine conditions. IC_50_ curves and values were generated with GraphPad Prism V10.0 using non-linear fit of log (inhibitor) versus response (three parameters).

### Western Blots

Cells washed with ice cold PBS were lysed in 1 x RIPA buffer supplemented with protease and phosphatase inhibitor cocktails I and II. Protein concentrations were determined by BCA kit on a microplate reader. Comparable amounts of protein (30μg) were resolved using NuPAGE 4%–12% Bis–Tris gradient gels and transferred to 0.2μm nitrocellulose membrane. After blocking with 5% BSA membranes were incubated overnight at 4°C with primary antibodies, followed by incubation with goat anti-rabbit or goat anti-mouse secondary antibodies coupled to horseradish peroxidase (HRP) in 5% nonfat milk for 1 hour at room temperature. Immunoblots were developed using enhanced chemiluminescence reagent and signal captured using the iBright CL1000 Imaging system. Reagents and antibodies are listed in **Supplementary Tables 3 and 6**.

### Statistics

Multivariable Cox regression models assessed the association between CH and OS. OS was measured from the time of the MSK-IMPACT testing blood draw to death or last follow-up. If the matched tumor sample was biopsied after the blood draw date, patients entered the model risk set at the time of biopsy using left truncation. The association between CH and OS was adjusted for age at the time of blood draw using a penalized B-spline in the Cox model. Kaplan-Meier survival probabilities were estimated for the categories of CH and similarly used left truncation. All survival analyses were conducted using R v4.4.1.

All the murine data presented was plotted and analyzed using GraphPad Prism 10, except for the clinical data and the CITEseq analysis, which is described in their figure legends or the CITEseq analysis section, respectively. Individual dots in the graphs represent distinct samples.

## Reporting summary

Further information on research design is available in the Reporting Summary linked to this article. All unique biological materials are available from the corresponding authors upon request or from the standard commercial sources described. Individual mouse strains presented in this study are available upon request or through the Jackson Laboratory.

## Data availability

The minimal data necessary to replicate the findings, except those in Figure 1 and Extended Data Figure 1, will be made publicly available. Clinical data cannot be made public to preserve patient anonymity. Additionally, raw sequencing data cannot be publicly deposited for legal and privacy reasons, as sequencing was performed for clinical purposes. Mutation calls will be available at the time of publication through cBioPortal. CITEseq data has been uploaded to GEO under accession number GSE275706.

## Code availability

All the code includes publicly available algorithms. All the code used for analysis for the CITEseq data will be available at the time of publication.

## Acknowledgements

We thank the members of the Fagin and Levine labs along with the friends and family of the main authors for their support and feedback. We acknowledge the generosity of patients and our early funders for this project including the Ludwig Center for Cancer Immunotherapy and the Cycle for Survival fund for rare cancer research at MSKCC (J.A.F, and R.L.L are co-PIs). V.T. is supported by the German Research Foundation (Deutsche Forschungsgemeinschaft DFG), the Center for Experimental Immuno-Oncology and the Mark Foundation for Cancer Research. P.S.V. is supported by the Mark Foundation for Cancer Research. R.S. is supported by NIH 1 RM1 HG011014-01. R.P.K. is supported in part by NIH/NCI P50 CA254838-01. J.A.F. is supported by ROI CA 255211 (PI), RO1 CA 249663 (J.A.F., Ho A, co-PIs). R.L.L is supported by the NIH NCI R35 (5R35CA197594) and the NCI Cancer Center Support Grant (CCSG, P30 CA08748).

We acknowledge the use of MSKCC’ Integrated Genomics Operation Core, Flow Cytometry Core Facility, Antitumour Assessment Core Facility, Animal Imaging Core Facility, Radiation Core Facility, Glassware Washing Core Facility, Media Preparation Core Facility, and the Research Animal Resource Center. The cores at MSKCC are funded by the NCI Cancer Center Support Grant (CCSG, P30 CA08748). Additionally, the Integrated Genomics Operation Core is funded by the Cycle for Survival, and the Marie-Josée and Henry R. Kravis Center for Molecular Oncology.

## Author information

These authors contributed equally: Vera Tiedje and Pablo Sánchez-Vela.

These authors jointly supervised this work: James A. Fagin and Ross L. Levine.

### Contributions

Conceptualization (Ideas; formulation or evolution of overarching research goals and aims): V.T., P.S.V., J.K., R.L.B., J.A.F, R.L.L.

Methodology (Development or design of methodology; creation of models): V.T, P.S.V, J.A.F., R.L.L.

Software (Programming, software development; designing computer programs; implementation of the computer code and supporting algorithms; testing of existing code components): J.L.Y., A.B.C., A.S., M.S.

Validation (Verification, whether as a part of the activity or separate, of the overall replication/ reproducibility of results/experiments and other research outputs): V.T., P.S.V., B.R.U., L.B.,

Formal analysis (Application of statistical, mathematical, computational, or other formal techniques to analyze or synthesize study data): V.T., P.S.V., J.L.Y., L.B., A.J.S., A.B.C., S.F.E., A.S., M.S., G.P.K., S.D.

Investigation (Conducting a research and investigation process, specifically performing the experiments, or data/evidence collection): V.T., P.S.V., B.R.U., L.B., A.J.S., A.B.C., S.F.E., J.G., M.W., A.K., K.C., T.Q., S.Y.I., A.K., A.R.M.B., R.P.

Resources (Provision of study materials, reagents, materials, patients, laboratory samples, animals, instrumentation, computing resources, or other analysis tools): A.J.S., S.F.E., M.K., E.D.S., A.Z., R.S., M.E., M.F.B., R.P.K., J.A.F., R.L.L.

Data Curation (Management activities to annotate (produce metadata), scrub data and maintain research data (including software code, where it is necessary for interpreting the data itself) for initial use and later reuse): V.T., P.S.V., J.L.Y., L.B., A.J.S., A.B.C., S.F.E., A.S., M.K., K.M., M.S., G.P.K., A.Z., R.S., S.D., M.E., M.F.B., R.P.K.,

Writing - Original Draft (Preparation, creation and/or presentation of the published work, specifically writing the initial draft including substantive translation): V.T, P.S.V, J.L.Y, R.P.K., J.A.F, R.L.L

Writing - Review & Editing (Preparation, creation and/or presentation of the published work by those from the original research group, specifically critical review, commentary or revision – including pre-or postpublication stages): V.T, P.S.V, J.A.F, R.L.L

Visualization (Preparation, creation and/or presentation of the published work, specifically visualization/ data presentation): V.T, P.S.V, J.L.Y., A.J.S, A.B.C, L.B.,, S.D.

Supervision (Oversight and leadership responsibility for the research activity planning and execution, including mentorship external to the core team): V.T., P.S.V., M.E., M.F.B., R.P.K., J.A.F, R.L.L

Project administration (Management and coordination responsibility for the research activity planning and execution): V.T, P.S.V, J.A.F, R.L.L

Funding acquisition (Acquisition of the financial support for the project leading to this publication): V.T, P.S.V, J.A.F., R.L.L.

## Ethics declaration

### Competing interests

V.T., P.S.V, J.L.Y., B.R.U., L.B., A.B.C., S.F.E., A.S., M.K., J.G, M.W., A.K., K.C., T.Q., A.R.M.B, R.P., M.S., G.P.K., E.D.S., A.Z., R.S., J.K., R.L.B, S.D. declare no relevant conflict of interest. A.J.S. declares spousal employment at Astra Zeneca. M.E. declares past grants from Ferrer International and Incyte, and personal fees from Quimatryx, unrelated to this work. M.F.B. declares consulting positions at AstraZeneca and Paige.AI and intellectual property rights at SOPHiA Genetics. R.P.K. is a co-founder of and consultant for Econic Biosciences. J.A.F. is a co-inventor of intellectual property focused on HRAS as a biomarker for treating cancer using tipifarnib which has been licensed by MSK to Kura Oncology. J.A.F. received prior research funding from Eisai and was a former consultant for LOXO Oncology, both unrelated to this work. R.L.L. is on the Supervisory board of Qiagen, a co-founder/advisor/board member at Ajax, and is a scientific advisor to Mission Bio, Zentalis, Auron, Prelude, Anovia, Bakx, Epiphanes, Kurome, Scorpion, Syndax and C4 Therapeutics; for each of these entities he receives equity. He has received research support from Calico, Zentalis and Ajax, and has consulted Jubilant, Goldman Sachs, Incyte, Astra Zeneca and Janssen.

### Ethics & Inclusion statement

We declare that the research included the local researchers throughout the research process – study design, study implementation, data ownership, intellectual property and authorship of publications. The research is locally relevant and has been determined in collaboration with local partners. The roles and responsibilities were agreed amongst collaborators ahead of the research and any capacity-building plans for local researchers were discussed. This study has been approved and complied with the regulations of the local ethics review committees for human research (IRB – Institutional Review Board), animal research (IACUC – Institutional Animal Care and Use Committee), and biosafety research practices (IBC – Institutional Biosafety Committee) at Memorial Sloan Kettering Cancer Center.

## Extended Data Figures and Tables

**Extended Data Fig. 2:**
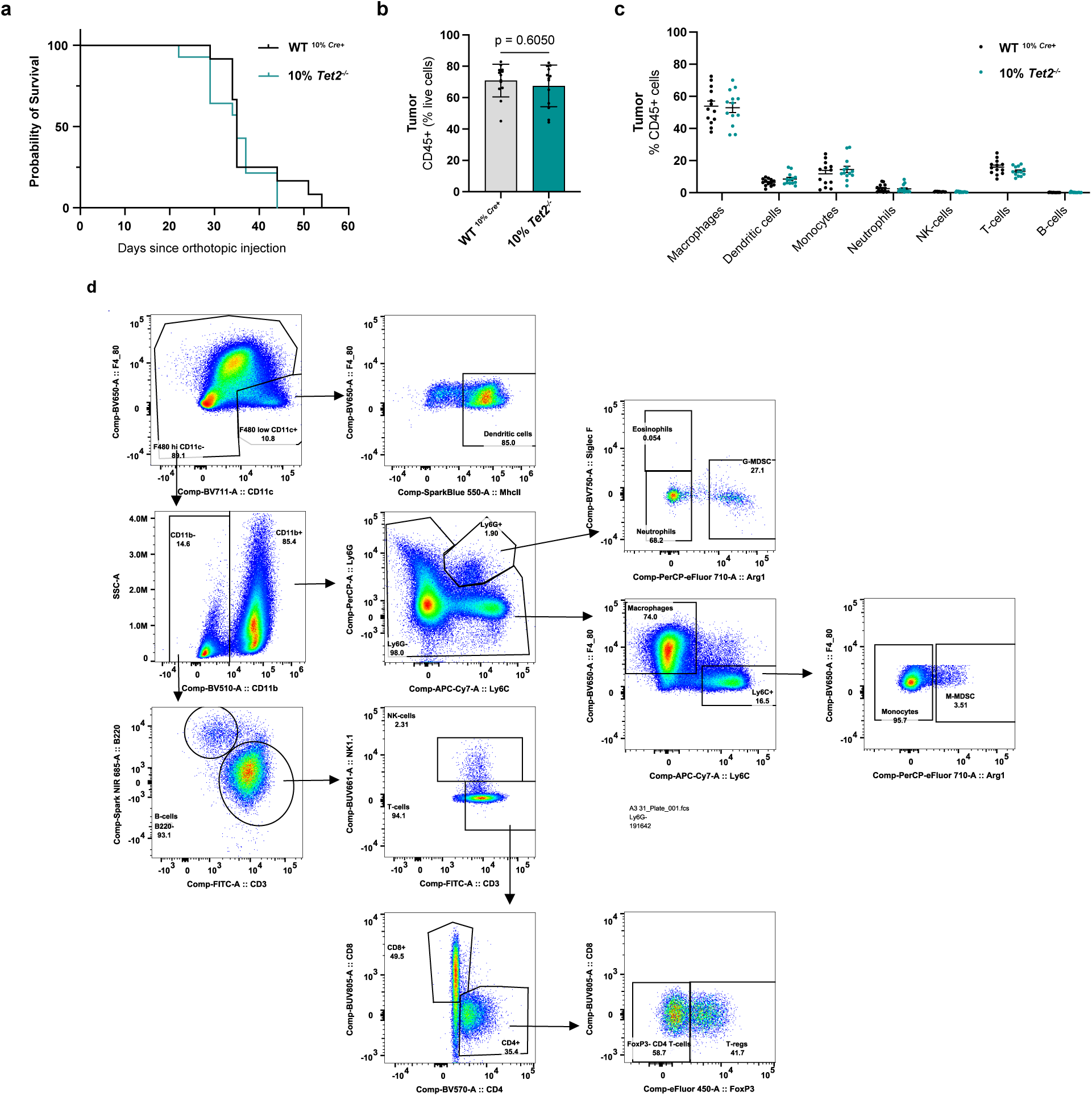
Baseline ATC-CH model characteristics and flow cytometric analysis. **a,** Kaplan-Meier curves of OS from tumour implantation date until humane-endpoints were met (development of early signs of tachypnea, weight loss, and overall decrease in activity) for 10% *Tet2^−/−^* (n=14) and WT ^10%^ *^Cre+^* (n=13) mice; **b**, Fraction of CD45+ cells of live cells in ATC tumours from WT ^10%^ *^Cre+^* and 10% *Tet2^−/−^* mice (n=13 each) **c**, Overall TME composition of ATC tumours from 13 10% *Tet2^−/−^* and WT ^10%^ *^Cre+^* mice each; **d**, Gating strategy in FlowJo Software for FACS analysis. Log-rank (Mantel-Cox) test (a); Bars represent mean with SEM, two-sided Multiple Mann Whitney U tests and Two-stage step-up method of Benjamini, Krieger and Yekutieli for FDR approach (b,c). OS: Overall survival; WT: Wild-type; ATC: Anaplastic thyroid cancer; TME: tumour microenvironment; FACS: Fluorescence-activated cell sorting; G-MDSC: Granulocytic-Myeloid Derived Suppressor Cells; NK-cells: Natural Killer cells; M-MDSC: Monocytic-Myeloid Derived Suppressor Cells. SEM: standar error of the mean.

**Extended Data Fig. 3:**
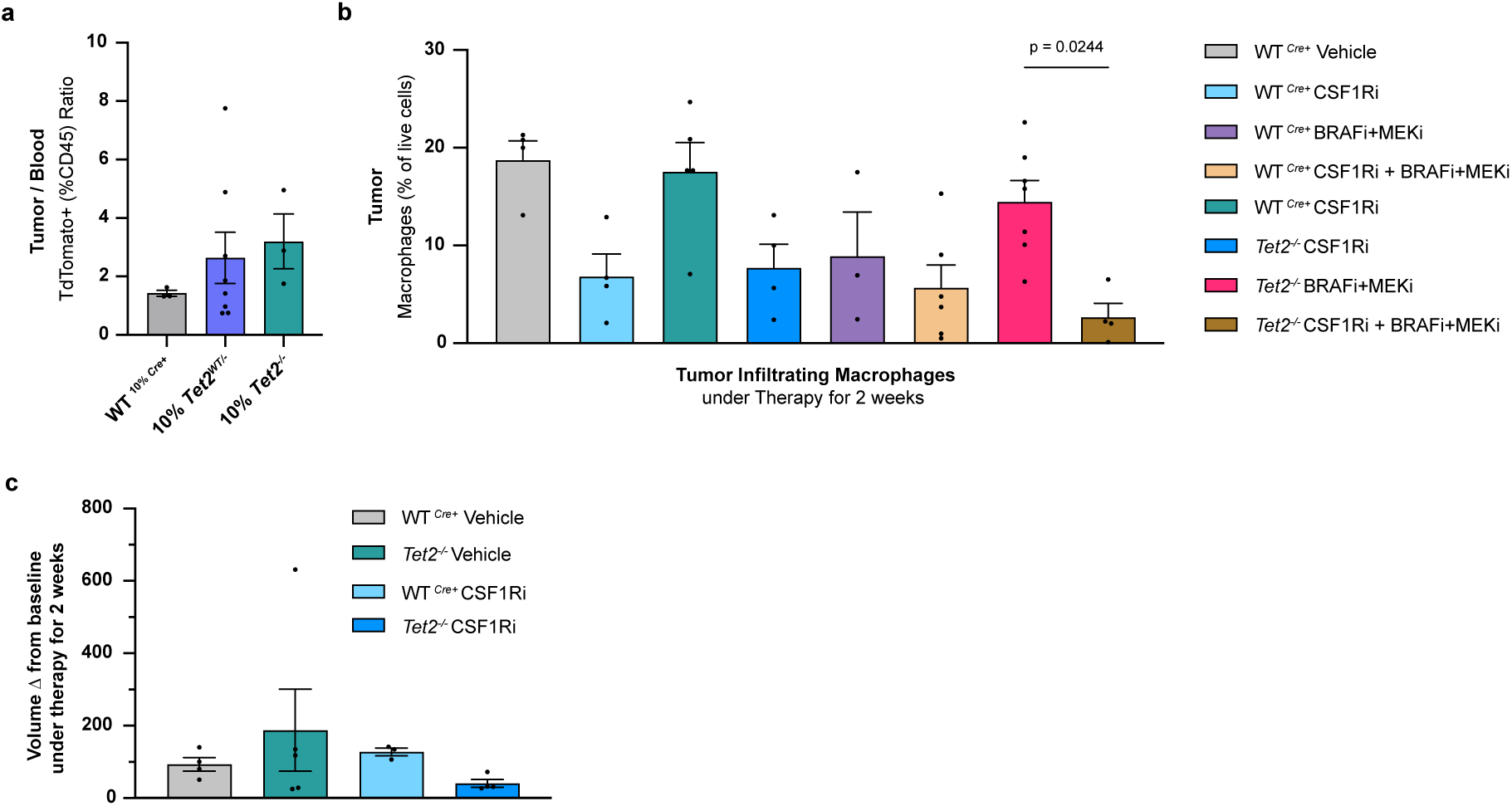
*Tet2^WT/-^* chimerism and macrophage depletion. **a,** Reporter chimerism ratio between tumour and blood for WT ^10%^ *^Cre+^* (n=3), 10% *Tet2^+/−^* (n=9) and 10% *Tet2^−/−^* (n=4); **b**, Tumour infiltrating macrophage (CD11b+ Ly6G-Ly6C-F480+) fraction of live cells from mice treated with vehicle (WT n=4, *Tet2^−/−^* n=5), CSF1Ri (WT n=4, *Tet2^−/−^* n=4), BRAFi+MEKi (WT n=3, *Tet2^−/−^* n=7) and BRAFi+MEKi in combination with CSF1Ri (WT n=4, *Tet2^−/−^* n=6) for 2 weeks; **c**, Change in tumour volume from baseline after 2 weeks of treatment (WT Vehicle n=4, WT CSF1Ri n=3, *Tet2^−/−^* Vehicle n=5 and *Tet2^−/−^* CSF1Ri n=4). Bars represent mean with SEM, two-sided Multiple Mann Whitney U tests and Two-stage step-up method of Benjamini, Krieger and Yekutieli for FDR approach (**b**). ATC: Anaplastic thyroid cancer; WT: Wild-type; CSF1Ri: Colony stimulating factor 1 receptor inhibitor; BRAFi: BRAF-inhibitor; MEKi: MEK-inhibitor. SEM: standard error of the mean.

**Extended Data Fig. 4.**
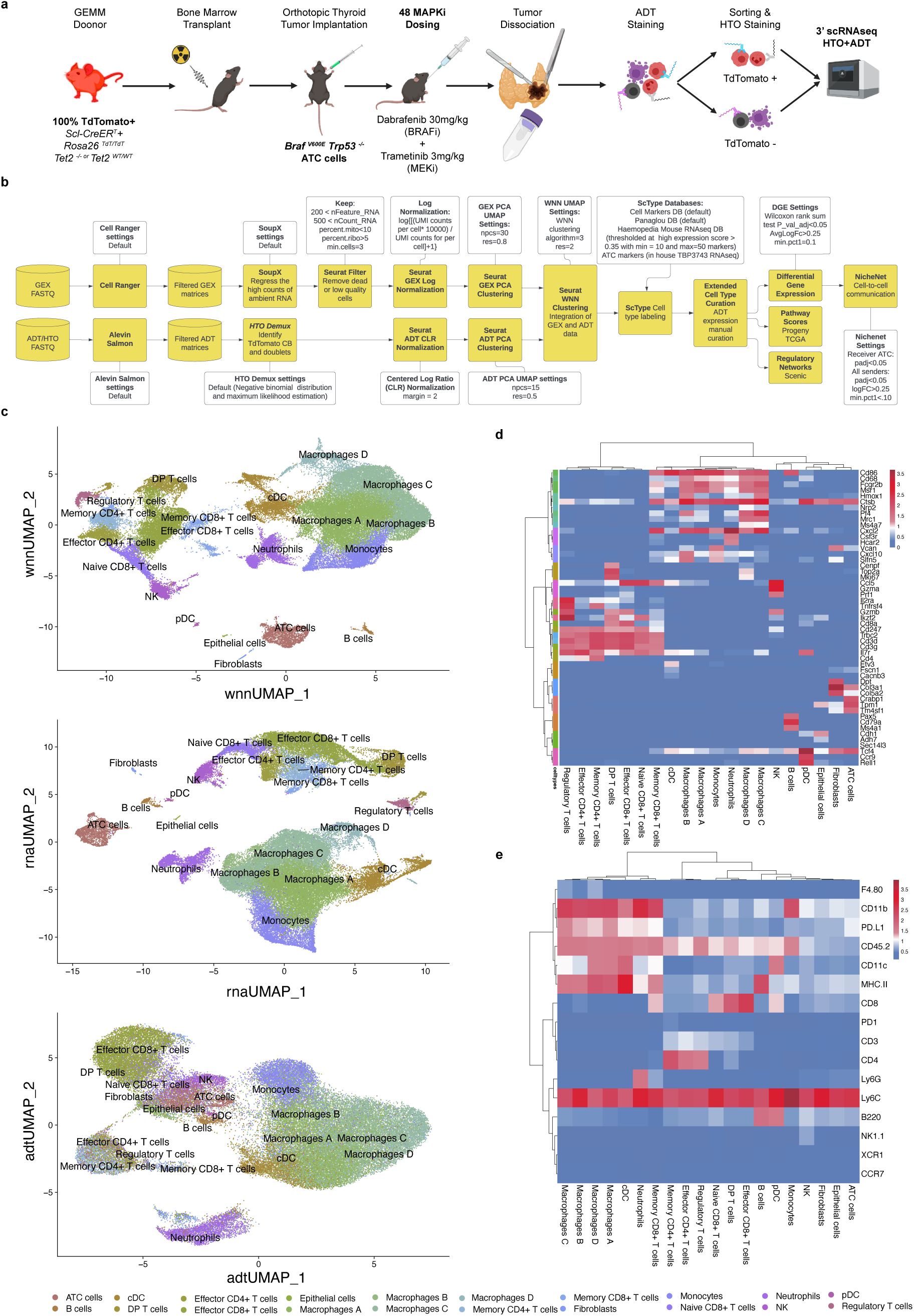
CITEseq of ATC tumours. **a,** Schema depicting design of the CITE-seq experiment; **b**, Flowchart showing the CITE-seq analysis steps. The different algorithms used are shown in yellow boxes. The parameters used at each step are shown in the white boxes. The intermediate data are shown in yellow cylinders; **c**, WNN, RNA, ADT UMAPs of single cell experiments. Weighted nearest neighbor WNN UMAP of the fully integrated data set with resolution (res = 2) and the UMAPs showing only the RNA data with number of pcs (npcs = 30) and resolution (res = 0.8) and ADT data with number of pcs (npcs = 15) and resolution (res =0.5); **d**, Heatmap of average RNA expression in cell populations. Hierarchical Ward D linkage clustering based on Pearson correlation of the average RNA expression within cell populations and shows the separation of the cell-types into larger subclusters and even the ordering of the cell-types with in their lymphoid and myeloid cell subclusters (i.e. CD4 and CD8 T cells cluster more closely together with themselves within the lymphoid populations and the Macrophage populations tend to cluster together within the myeloid subcluster). **e**, Heatmap of average ADT expression in cell populations. Hierarchical clustering of the average ADT expression with a Ward D linkage and Pearson correlation metric within cell populations and show the separation of the lymphoid and myeloid cell types as well a clear separation of the infiltrating immune cells from the tumour and nearby tissue cell types.

**Extended Data Fig. 5:**
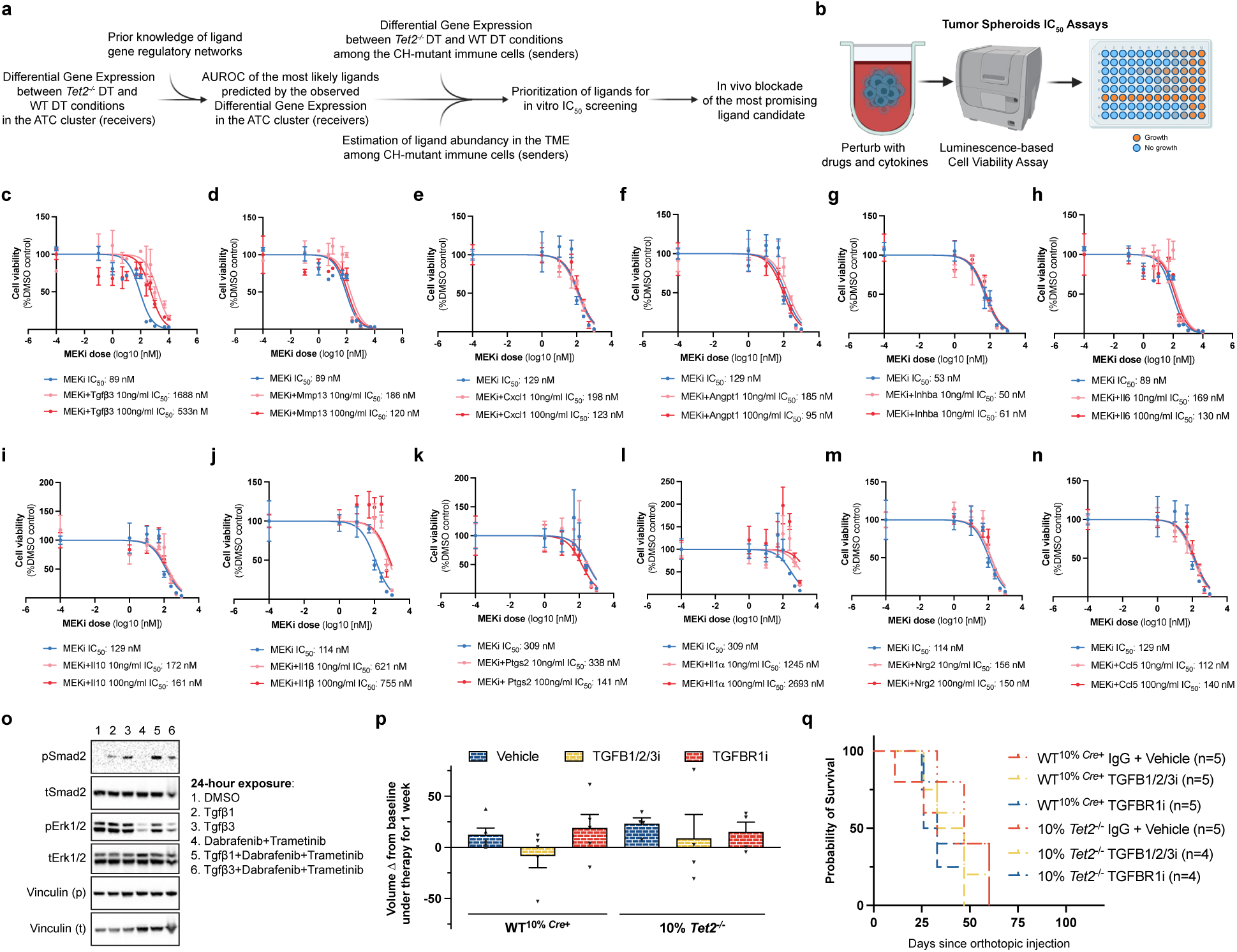
Mechanism workup. **a**, Proposed hypothesis-testing validating workflow; **b**, Tumour spheroids assay set up for ligand screening; **c-n**, IC50s of non-validating ligands; **o**, Western blot of TBP3743 ATC cells treated *in vitro* for 24 h with 1nM dabrafenib (BRAFi) and trametinib (MEKi), respectively, probing for pERK, tERK, pSMAD2, tSMAD2 and Vinculin; **p**, Tumour volume after one week of treatment with vehicle/IgG (WT ^10%^ *^Cre+^* n=5, 10% *Tet2^−/−^* n=4), TGFBR1i (WT ^10%^ *^Cre+^* n=5 and 10% *Tet2^−/−^* n= 4) or TGFB1/2/3i (WT ^10%^ *^Cre+^* n=5, 10% *Tet2^−/−^* n=4) ; q, OS from tumour implantation date from the vehicle/IgG WT ^10%^ *^Cre+^* n=5, 10% *Tet2^−/−^* n=5), TGFBR1i (WT ^10%^ *^Cre+^* n=5, 10% *Tet2^−/−^*n=4) or TGFB1/2/3i (WT ^10%^ *^Cre+^* n=5, 10% *Tet2^−/−^* n=4). Bars represent mean with SEM. Two-sided Multiple Mann Whitney U tests and Two-stage step-up method of Benjamini, Krieger and Yekutieli for FDR approach (**p**); Log-rank (Mantel-Cox) test (**q**). BRAFi: BRAF inhibitor; MEKi: MEK inhibitor; WT: Wildt-type; TGFBR1i: TGFB receptor 1 inhibitor; TGFB1/2/3i: TGFB 1/2/3 inhibitor. OS: overall survival. SEM: Standard error of the mean.

**Extended Table 1:** ATC cohort characteristics.

| **Characteristic** | **N = 89<sup>1</sup>** |
| --- | --- |
| **Age at MSK-IMPACT (blood)** | 66 (33, 88) |
| **Age group** |  |
| (0,55) | 15 (17%) |
| (55,65) | 25 (28%) |
| (65,75) | 28 (31%) |
| (75,100) | 21 (24%) |
| **Sex** |  |
| Female | 39 (44%) |
| Male | 50 (56%) |
| **Smoking History** | 37 (42%) |
| **Prior Therapy Received** | 30 (34%) |
| **Any MAPK inhibitor Therapy Received** | 11 (12%) |
| **Dabrafenib-Trametinib Received** | 7 (7.9%) |
| **ATC Stage** |  |
| Metastatic (Stage IVc) | 50 (56%) |
| Non-metastatic (Stage IVa-b) | 39 (44%) |
| **ATC Type** |  |
| Primary | 70 (79%) |
| Transformed from DTC | 19 (21%) |
| **CH classification by variant** |  |
| No CH | 56 (63%) |
| CH-PD | 20 (22%) |
| CH non-PD | 13 (15%) |
| **CH classification by gene (CH-PD)** |  |
| No CH | 56 (81%) |
| *DNMT3A* PD | 9 (13%) |
| *TET2* PD | 4 (5.8%) |
| Other | 20 |
| **CH classification by gene (All CH)** |  |
| No CH | 56 (80%) |
| *DNMT3A* CH | 10 (14%) |
| *TET2* CH | 4 (5.7%) |
| Other | 19 |
| **CH classification by high blood VAF (nVAF >0.2)** |  |
| No CH | 56 (86%) |
| High nVAF CH non-PD | 3 (4.6%) |
| High nVAF CH-PD | 6 (9.2%) |
| Unknown | 24 |
| **CH classification by high tumor VAF (tVAF >0)** |  |
| No CH | 56 (63%) |
| High tVAF CH non-PD | 14 (16%) |
| High tVAF CH-PD | 19 (21%) |

<sup>1</sup>Median (Min, Max); n (%)

**Extended Table 2:** Variables associated with the presence of CH.

| **Characteristic** | **Overall**<br>**N = 89<sup>1</sup>** | **No CH**<br>**N = 56<sup>1</sup>** | **CH-PD**<br>**N = 20<sup>1</sup>** | **CH non-PD**<br>**N = 13<sup>1</sup>** | **p-value<sup>2</sup>** |
| --- | --- | --- | --- | --- | --- |
| **Age at MSK-IMPACT (blood)** | 66 (33, 88) | 63 (33, 85) | 71 (45, 88) | 70 (44, 86) | 0.025 |
| **Age group** |  |  |  |  | 0.079 |
| (0,55) | 15 (17%) | 12 (21%) | 2 (10%) | 1 (7.7%) |  |
| (55,65) | 25 (28%) | 20 (36%) | 4 (20%) | 1 (7.7%) |  |
| (65,75) | 28 (31%) | 13 (23%) | 7 (35%) | 8 (62%) |  |
| (75,100) | 21 (24%) | 11 (20%) | 7 (35%) | 3 (23%) |  |
| **Sex** |  |  |  |  | 0.5 |
| Female | 39 (44%) | 22 (39%) | 10 (50%) | 7 (54%) |  |
| Male | 50 (56%) | 34 (61%) | 10 (50%) | 6 (46%) |  |
| **Smoking History** | 37 (42%) | 20 (36%) | 9 (45%) | 8 (62%) | 0.2 |
| **Prior Therapy Received** | 30 (34%) | 19 (34%) | 7 (35%) | 4 (31%) | >0.9 |
| **Any MAPK inhibitor Therapy Received** | 11 (12%) | 8 (14%) | 2 (10%) | 1 (7.7%) | >0.9 |
| **Dabrafenib-Trametinib Received** | 7 (7.9%) | 5 (8.9%) | 1 (5.0%) | 1 (7.7%) | >0.9 |
| **ATC Stage** |  |  |  |  | 0.8 |
| Metastatic (Stage IVc) | 50 (56%) | 30 (54%) | 12 (60%) | 8 (62%) |  |
| Non-metastatic (Stage IVa-b) | 39 (44%) | 26 (46%) | 8 (40%) | 5 (38%) |  |
| **ATC Type** |  |  |  |  | >0.9 |
| Primary | 70 (79%) | 44 (79%) | 16 (80%) | 10 (77%) |  |
| Transformed from DTC | 19 (21%) | 12 (21%) | 4 (20%) | 3 (23%) |  |

<sup>1</sup>Median (Min, Max); n (%)
<sup>2</sup>Kruskal-Wallis rank sum test; Fisher's exact test; Pearson's Chi-squared test

**Extended Table 3:** Univariable OS Regression.

| **Characteristic** | **Total N** | **Event N** | **HR<sup>1</sup>** | **95% CI<sup>1</sup>** | **p-value** |
| --- | --- | --- | --- | --- | --- |
| **Age at MSK-IMPACT (blood)** | 89 | 74 | 1.01 | 0.99, 1.03 | 0.3 |
| **Sex** | 89 | 74 |  |  |  |
| Female |  |  | — | — |  |
| Male |  |  | 0.67 | 0.42, 1.08 | 0.10 |
| **Smoking History** | 89 | 74 | 0.99 | 0.62, 1.57 | >0.9 |
| **Prior Therapy Received** | 89 | 74 | 1.24 | 0.76, 2.03 | 0.4 |
| **Any MAPK inhibitor Therapy Received** | 89 | 74 | 0.77 | 0.38, 1.55 | 0.5 |
| **Dabrafenib-Trametinib Received** | 89 | 74 |  |  |  |
| No |  |  | — | — |  |
| Yes |  |  | 0.64 | 0.26, 1.60 | 0.3 |
| **ATC Stage** | 89 | 74 |  |  |  |
| Metastatic (Stage IVc) |  |  | — | — |  |
| Non-metastatic (Stage IVa-b) |  |  | 0.31 | 0.18, 0.52 | <0.001 |
| **ATC Type** | 89 | 74 |  |  |  |
| Primary |  |  | — | — |  |
| Transformed from DTC |  |  | 1.28 | 0.72, 2.26 | 0.4 |
| **CH classification by variant** | 89 | 74 |  |  |  |
| No CH |  |  | — | — |  |
| CH-PD |  |  | 2.68 | 1.49, 4.81 | <0.001 |
| CH non-PD |  |  | 1.74 | 0.89, 3.40 | 0.11 |
| **CH classification by variant (CH-PD only)** | 76 | 63 |  |  |  |
| No CH |  |  | — | — |  |
| CH-PD |  |  | 2.75 | 1.51, 4.98 | <0.001 |
| **CH classification by high blood VAF (nVAF >0.2)** | 65 | 52 |  |  |  |
| No CH |  |  | — | — |  |
| High nVAF CH non-PD |  |  | 0.71 | 0.17, 2.95 | 0.6 |
| High nVAF CH-PD |  |  | 1.87 | 0.73, 4.82 | 0.2 |
| **CH classification by high tumor VAF (tVAF>0)** | 89 | 74 |  |  |  |
| No CH |  |  | — | — |  |
| High tVAF CH non-PD |  |  | 1.82 | 0.95, 3.48 | 0.071 |

| Characteristic | Total N | Event N | HR<sup>1</sup> | 95% CI<sup>1</sup> | p-value |
| --- | --- | --- | --- | --- | --- |
| High tVAF CH-PD |  |  | 2.63 | 1.45, 4.76 | 0.001 |

<sup>1</sup>HR = Hazard Ratio, CI = Confidence Interval

**Extended Table 4:** Multivariate OS Regression Adjusted for Age using a Spline.

| **Characteristic<sup>1</sup>** | **Event N<sup>1</sup>** | **HR<sup>12</sup>** | **95% CI<sup>12</sup>** | **p-value<sup>1</sup>** |
| --- | --- | --- | --- | --- |
| **CH classification by variant** |  |  |  |  |
| No CH | 45 | — | — |  |
| CH-PD | 18 | 2.84 | 1.51, 5.36 | 0.001 |
| CH non-PD | 11 | 1.79 | 0.88, 3.68 | 0.11 |

<sup>1</sup>Adjusts for age using spline
<sup>2</sup>HR = Hazard Ratio, CI = Confidence Interval

| **Characteristic<sup>1</sup>** | **Event N<sup>1</sup>** | **HR<sup>12</sup>** | **95% CI<sup>12</sup>** | **p-value<sup>1</sup>** |
| --- | --- | --- | --- | --- |
| **CH classification by variant (CH-PD only)** |  |  |  |  |
| No CH | 45 | — | — |  |
| CH-PD | 18 | 2.87 | 1.50, 5.48 | 0.001 |

<sup>1</sup>Adjusts for age using spline
<sup>2</sup>HR = Hazard Ratio, CI = Confidence Interval

**Extended Table 5:** Multivariate OS Regression Adjusted for Metastatic Disease and Age using a Spline.

| Characteristic<sup>1</sup> | Event N<sup>1</sup> | HR<sup>12</sup> | 95% CI<sup>12</sup> | p-value<sup>1</sup> |
| --- | --- | --- | --- | --- |
| **CH classification by variant** |  |  |  |  |
| No CH | 45 | — | — |  |
| CH-PD | 18 | 2.16 | 1.12, 4.14 | 0.021 |
| CH non-PD | 11 | 1.40 | 0.68, 2.88 | 0.4 |
| **ATC Stage** |  |  |  |  |
| Metastatic (Stage IVc) | 48 | — | — |  |
| Non-metastatic (Stage IVa-b) | 26 | 0.32 | 0.18, 0.55 | <0.001 |

<sup>1</sup>Adjusts for age using spline
<sup>2</sup>HR = Hazard Ratio, CI = Confidence Interval

| Characteristic<sup>1</sup> | Event N<sup>1</sup> | HR<sup>12</sup> | 95% CI<sup>12</sup> | p-value<sup>1</sup> |
| --- | --- | --- | --- | --- |
| **CH classification by variant (CH-PD only)** |  |  |  |  |
| No CH | 45 | — | — |  |
| CH-PD | 18 | 2.20 | 1.13, 4.28 | 0.020 |
| **ATC Stage** |  |  |  |  |
| Metastatic (Stage IVc) | 40 | — | — |  |
| Non-metastatic (Stage IVa-b) | 23 | 0.30 | 0.16, 0.54 | <0.001 |

<sup>1</sup>Adjusts for age using spline
<sup>2</sup>HR = Hazard Ratio, CI = Confidence Interval

## Supplementary information (e.g.: Reporting summary and supplementary tables)

Reporting Summary and Editorial Checklist have been provided.

## Source data

Supplementary tables with the raw data to facilitate reproducibility of the main non-clinical findings will be provided at the time of publication.

## References

1. Molinaro, E. et al. Anaplastic thyroid carcinoma: from clinicopathology to genetics and advanced therapies. Nat Rev Endocrinol 13, 644–660 (2017).

2. Tiedje, V. et al. NGS based identification of mutational hotspots for targeted therapy in anaplastic thyroid carcinoma. Oncotarget 8, 42613–42620 (2017).

3. Pozdeyev, N. et al. Genetic Analysis of 779 Advanced Differentiated and Anaplastic Thyroid Cancers. Clinical Cancer Research 24, 3059–3068 (2018).

4. Landa, I. et al. Genomic and transcriptomic hallmarks of poorly differentiated and anaplastic thyroid cancers. J Clin Invest 126, 1052–1066 (2016).

5. Ryder, M., Ghossein, R. A., Ricarte-Filho, J. C. M., Knauf, J. A. & Fagin, J. A. Increased density of tumor-associated macrophages is associated with decreased survival in advanced thyroid cancer. Endocr Relat Cancer 15, 1069–1074 (2008).

6. Luo, H. et al. Characterizing dedifferentiation of thyroid cancer by integrated analysis. Sci Adv 7, eabf3657 (2024).

7. Lu, L. et al. Anaplastic transformation in thyroid cancer revealed by single-cell transcriptomics. J Clin Invest 133, (2023).

8. Subbiah, V. et al. Dabrafenib and Trametinib Treatment in Patients With Locally Advanced or Metastatic BRAF V600–Mutant Anaplastic Thyroid Cancer. Journal of Clinical Oncology 36, 7–13 (2017).

9. Maniakas, A. et al. Evaluation of Overall Survival in Patients With Anaplastic Thyroid Carcinoma, 2000-2019. JAMA Oncol 6, 1397–1404 (2020).

10. Subbiah, V. et al. Dabrafenib plus trametinib in patients with BRAF V600E-mutant anaplastic thyroid cancer: updated analysis from the phase II ROAR basket study. Annals of Oncology 33, 406–415 (2022).

11. Busque, L. et al. Recurrent somatic TET2 mutations in normal elderly individuals with clonal hematopoiesis. Nat Genet 44, 1179–1181 (2012).

12. Coombs, C. C. et al. Therapy-Related Clonal Hematopoiesis in Patients with Non-hematologic Cancers Is Common and Associated with Adverse Clinical Outcomes. Cell Stem Cell 21, 374–382.e4 (2017).

13. Bolton, K. L. et al. Cancer therapy shapes the fitness landscape of clonal hematopoiesis. Nat Genet 52, 1219–1226 (2020).

14. Boucai, L. et al. Radioactive Iodine–Related Clonal Hematopoiesis in Thyroid Cancer Is Common and Associated With Decreased Survival. J Clin Endocrinol Metab 103, 4216–4223 (2018).

15. Moran-Crusio, K. et al. Tet2 Loss Leads to Increased Hematopoietic Stem Cell Self-Renewal and Myeloid Transformation. Cancer Cell 20, 11–24 (2011).

16. Fuster, J. J. et al. Clonal hematopoiesis associated with TET2 deficiency accelerates atherosclerosis development in mice. Science (1979) 355, 842–847 (2017).

17. Sánchez Vela, P., Trowbridge, J. J. & Levine, R. L. Clonal hematopoiesis, aging and Alzheimer’s disease. Nat Med 29, 1605–1606 (2023).

18. Ptashkin, R. N. et al. Prevalence of Clonal Hematopoiesis Mutations in Tumor-Only Clinical Genomic Profiling of Solid Tumors. JAMA Oncol 4, 1589–1593 (2018).

19. Buscarlet, M. et al. Lineage restriction analyses in CHIP indicate myeloid bias for TET2 and multipotent stem cell origin for DNMT3A. Blood 132, 277–280 (2018).

20. Göthert, J. R. et al. In vivo fate-tracing studies using the Scl stem cell enhancer: embryonic hematopoietic stem cells significantly contribute to adult hematopoiesis. Blood 105, 2724–2732 (2005).

21. Madisen, L. et al. A robust and high-throughput Cre reporting and characterization system for the whole mouse brain. Nat Neurosci 13, 133–140 (2010).

22. Vanden Borre, P., et al. The Next Generation of Orthotopic Thyroid Cancer Models: Immunocompetent Orthotopic Mouse Models of BRAFV600E-Positive Papillary and Anaplastic Thyroid Carcinoma. Thyroid® 24, 705–714 (2013).

23. Pyonteck, S. M. et al. CSF-1R inhibition alters macrophage polarization and blocks glioma progression. Nat Med 19, 1264–1272 (2013).

24. Stoeckius, M. et al. Simultaneous epitope and transcriptome measurement in single cells. Nat Methods 14, 865–868 (2017).

25. Agrawal, N. et al. Integrated Genomic Characterization of Papillary Thyroid Carcinoma. Cell 159, 676–690 (2014).

26. Schubert, M. et al. Perturbation-response genes reveal signaling footprints in cancer gene expression. Nat Commun 9, 20 (2018).

27. Browaeys, R., Saelens, W. & Saeys, Y. NicheNet: modeling intercellular communication by linking ligands to target genes. Nat Methods 17, 159–162 (2020).

28. Lee, M. K. et al. TGF-β activates Erk MAP kinase signalling through direct phosphorylation of ShcA. EMBO J 26, 3957–3967–3967 (2007).

29. Jin, C. H. et al. Discovery of N-((4-([1,2,4]Triazolo[1,5-a]pyridin-6-yl)-5-(6-methylpyridin-2-yl)-1H-imidazol-2-yl)methyl)-2-fluoroaniline (EW-7197): A Highly Potent, Selective, and Orally Bioavailable Inhibitor of TGF-β Type I Receptor Kinase as Cancer Immunotherapeutic/Antifibrotic Agent. J Med Chem 57, 4213–4238 (2014).

30. Dasch, J. R., Pace, D. R., Waegell, W., Inenaga, D. & Ellingsworth, L. Monoclonal antibodies recognizing transforming growth factor-beta. Bioactivity neutralization and transforming growth factor beta 2 affinity purification. The Journal of Immunology 142, 1536–1541 (1989).

31. Hanahan, D. & Weinberg, R. A. Hallmarks of Cancer: The Next Generation. Cell 144, 646–674 (2011).

32. Kessler, M. D. et al. Common and rare variant associations with clonal haematopoiesis phenotypes. Nature 612, 301–309 (2022).

33. Ruark, E. et al. Mosaic PPM1D mutations are associated with predisposition to breast and ovarian cancer. Nature 493, 406–410 (2013).

34. Feng, Y. et al. Hematopoietic-specific heterozygous loss of Dnmt3a exacerbates colitis-associated colon cancer. Journal of Experimental Medicine 220, e20230011 (2023).

35. Nguyen, Y. T. M. et al. Tet2 deficiency in immune cells exacerbates tumor progression by increasing angiogenesis in a lung cancer model. Cancer Sci 112, 4931–4943 (2021).

36. Narod, S. Response. JNCI: Journal of the National Cancer Institute 106, dju046 (2014).

37. Li, S. et al. TET2 promotes anti-tumor immunity by governing G-MDSCs and CD8-T-cell numbers. EMBO Rep 21, e49425 (2020).

38. Pan, W. et al. The DNA Methylcytosine Dioxygenase Tet2 Sustains Immunosuppressive Function of Tumor-Infiltrating Myeloid Cells to Promote Melanoma Progression. Immunity 47, 284–297.e5 (2017).

39. Jaiswal, S. et al. Clonal Hematopoiesis and Risk of Atherosclerotic Cardiovascular Disease. New England Journal of Medicine 377, 111–121 (2017).

40. Kim, P. G. et al. Dnmt3a-mutated clonal hematopoiesis promotes osteoporosis. Journal of Experimental Medicine 218, e20211872 (2021).

41. Agrawal, M. et al. TET2-mutant clonal hematopoiesis and risk of gout. Blood 140, 1094–1103 (2022).

42. Miller, P. G. et al. Association of clonal hematopoiesis with chronic obstructive pulmonary disease. Blood 139, 357–368 (2022).

43. Wong, W. J. et al. Clonal haematopoiesis and risk of chronic liver disease. Nature 616, 747–754 (2023).

44. Vlasschaert, C. et al. Clonal hematopoiesis of indeterminate potential is associated with acute kidney injury. Nat Med 30, 810–817 (2024).

45. Quin, C. et al. Neutrophil-mediated innate immune resistance to bacterial pneumonia is dependent on Tet2 function. J Clin Invest 134, (2024).

46. Huang, S. et al. MED12 Controls the Response to Multiple Cancer Drugs through Regulation of TGF-β Receptor Signaling. Cell 151, 937–950 (2012).

47. Cheng, D. T. et al. Memorial Sloan Kettering-Integrated Mutation Profiling of Actionable Cancer Targets (MSK-IMPACT): A Hybridization Capture-Based Next-Generation Sequencing Clinical Assay for Solid Tumor Molecular Oncology. The Journal of Molecular Diagnostics 17, 251–264 (2015).

48. Shen, R. & Seshan, V. E. FACETS: allele-specific copy number and clonal heterogeneity analysis tool for high-throughput DNA sequencing. Nucleic Acids Res 44, e131–e131 (2016).

49. Shen, F. W. et al. Cloning of Ly-5 cDNA. Proceedings of the National Academy of Sciences 82, 7360–7363 (1985).

50. Pham, T. H. et al. Machine-Learning and Chemicogenomics Approach Defines and Predicts Cross-Talk of Hippo and MAPK Pathways. Cancer Discov 11, 778–793 (2021).

